# Bitter taste genetics and oral health in Canadian Longitudinal Study on Aging

**DOI:** 10.1101/2024.11.12.623243

**Authors:** Marziyeh Shafizadeh, Vikram Bhatia, Samah Ahmed, Britt Drögemöller, Chrysi Stavropoulou, Philip St. John, Rajinder P. Bhullar, Prashen Chelikani, Carol A. Hitchon

**Affiliations:** Department of Oral Biology and Manitoba Chemosensory Biology Research Group, Dr. Gerald Niznick College of Dentistry, Rady Faculty of Health Sciences, University of Manitoba, Winnipeg, MB, Canada; Children’s Hospital Research Institute of Manitoba (CHRIM), University of Manitoba, Winnipeg, MB, Canada; Department of Biochemistry and Medical Genetics, Max Rady College of Medicine, Rady Faculty of Health Sciences, University of Manitoba, Winnipeg, MB, Canada; Paul Albrechtsen Research Institute CancerCare Manitoba, Winnipeg, MB, Canada; Centre on Aging, University of Manitoba, Winnipeg, MB, Canada; Department of Dental Diagnostic and Surgical Sciences, Dr. Gerald Niznick College of Dentistry, Rady Faculty of Health Sciences, University of Manitoba, Winnipeg, MB, Canada; Department of Internal Medicine, Max Rady College of Medicine, Rady Faculty of Health Sciences, University of Manitoba, Winnipeg, MB, Canada

**Keywords:** Taste receptor, taste genetics, pseudogene, oral health, CLSA, allelic association, single nucleotide polymorphism

## Abstract

This study aimed to investigate the association of single nucleotide polymorphisms (SNPs) in 25 Bitter Taste Receptor genes (*TAS2R*s) and 12 *TAS2R* pseudogenes with self-reported oral health outcomes in the Canadian Longitudinal Study on Aging (CLSA) cohort. Following quality control, 124 SNPs with a minor allele frequency > 0.01 and 21,991 individuals of European ancestry were included in the analysis. Fifteen SNPs in *TAS2R8, 9, 13, 14, 20*, and *50* were significantly associated with self-reported sore jaw muscles, a symptom commonly linked to temporomandibular disorders (TMDs). *TAS2R20* exhibited the highest number of associated SNPs. Structure-function analysis suggests that variants in *TAS2R20* may contribute to this symptom by altering ligand interactions. These findings highlight the potential for *TAS2R* genetic screening to identify individuals at elevated risk for TMD, supporting the development of personalized treatment strategies and advancing our understanding of TMD genetic risk factors.

## INTRODUCTION

Bitter Taste Receptors (T2Rs) are G-protein coupled receptors that detect bitter compounds. These receptors are encoded by genes approximately 1000 base pairs in length, referred to as *TAS2R* according to the HUGO Gene Nomenclature.^1^ In humans, there are 25 functional *TAS2R* genes and 12 pseudogenes.^2-6^ Pseudogenes are sequences that are similar to genes but cannot produce functional proteins due to genetic alterations, such as premature stop codons or frameshifts. Despite lacking protein-coding ability, growing evidence indicates that pseudogenes can still be transcribed and may play important regulatory roles in gene expression.^7-9^ *TAS2R*s exhibit various single nucleotide polymorphisms (SNPs), which can impact receptor expression and function.^10,11^

T2Rs play a crucial role in the innate immune response to oral pathogens. These receptors are expressed in various tissues, including the mucosal epithelium and immune cells.^12-14^ T2Rs activate pathways that lead to the release of nitric oxide and calcium-mediated antimicrobial peptides, which have direct antimicrobial effects.^15-17^ Studies have shown that genetic variants in *TAS2R*s can influence the composition of oral microbial communities.^18,19^ Variations in *TAS2R*s may alter receptor expression or function, impacting the immune response to oral microorganisms and, consequently, susceptibility to oral diseases.

Additionally, T2Rs can influence oral health through various mechanisms. These receptors affect food preferences and dietary habits, which are closely related to oral health issues such as dental caries and periodontal disease.^20-23^ A previous transcriptome-wide association study identified a significant relationship between the expression of certain T2Rs and the occurrence of early childhood caries.^24^ Furthermore, variants in *TAS2R*s are linked to alcohol consumption and smoking, which are known risk factors for periodontal diseases.^25-27^ A previous study demonstrated that individuals with specific genotypes of *TAS2R*38 have a lower risk of periodontitis.^28^

Few studies are available on the relationship between *TAS2R* genetic variants and oral health conditions. Available studies focused primarily on dental caries, while other conditions remain largely unexplored.^18,24,28-31^ Most of these studies have concentrated on the *TAS2R*38 variants.^28-31^ Research on the association between dental caries and other *TAS2R* types is limited and has yielded conflicting results.^18,24^ Furthermore, the association between other oral health problems and *TAS2R* variants remains poorly understood.

The Canadian Longitudinal Study on Aging (CLSA) is a cohort study of 50,000 individuals aged 45-85 years, of whom 26,622 underwent genome-wide genotyping of DNA samples collected from blood.^32,33^ Extensive demographic and clinical data on various aspects of participants’ lives have been collected to assess their impact on health and disease development.^33^ In this study, we analyzed the allele distributions of *TAS2R* SNPs in the CLSA dataset. Additionally, we evaluated the association between *TAS2R* SNPs and various self-reported oral health symptoms within the cohort. To understand the potential mechanisms underlying this association, we conducted a structure-function analysis of *TAS2R20* variants, focusing on their possible impact on ligand interactions.

## RESULTS

### Allele distributions of *TAS2R* SNPs in CLSA and 1KGP

A total of 1,171 non-monomorphic variants occurring within *TAS2R* genes and pseudogenes were extracted from the CLSA. Following the application of quality control criteria, 124 SNPs and 22,974 individuals of European ancestry were included in the allele distribution analyses. Of these SNPs, 87 were located in *TAS2R* genes and 37 in *TAS2R* pseudogenes. Detailed annotations are provided in Table S1. Notably, *TAS2R20* had the highest number of SNPs (n=15), followed by *TAS2R4* (n=12), *TAS2R42* (n=8), *TAS2R14* (n=7), and *TAS2R15P* (n=7). The distribution of functional consequences was as follows: missense (n=38), 3’ untranslated region (UTR) (n=20), synonymous (n=19), 5’ UTR (n=9), and stop-gained (n=1; in *TAS2R19*).

Allele frequencies for 121 out of 124 SNPs identified in the CLSA dataset were available for the European population in 1KGP. Minor allele frequencies in CLSA and the p-values from the chi-squared tests comparing them with the corresponding allele frequencies in the 1KGP European population are illustrated in **Figure 1** with details provided in Table S1. Chi-squared analysis revealed significant differences in allele frequency for 48 out of 121 SNPs between the two databases. Among these, 15 were missense variants, 14 were located in pseudogenes, 7 in untranslated regions, and 12 were synonymous variants. The highest proportion of variants with significantly different allele distributions was observed in *TAS2R15P*, where all seven variants exhibited differences (Table S2). *TAS2R20* also showed a substantial proportion, with significant differences in 11 out of 15 variants (73%). Pairwise linkage disequilibrium between the variants is shown in heatmaps in **Figure S1**.

**Figure 1.**
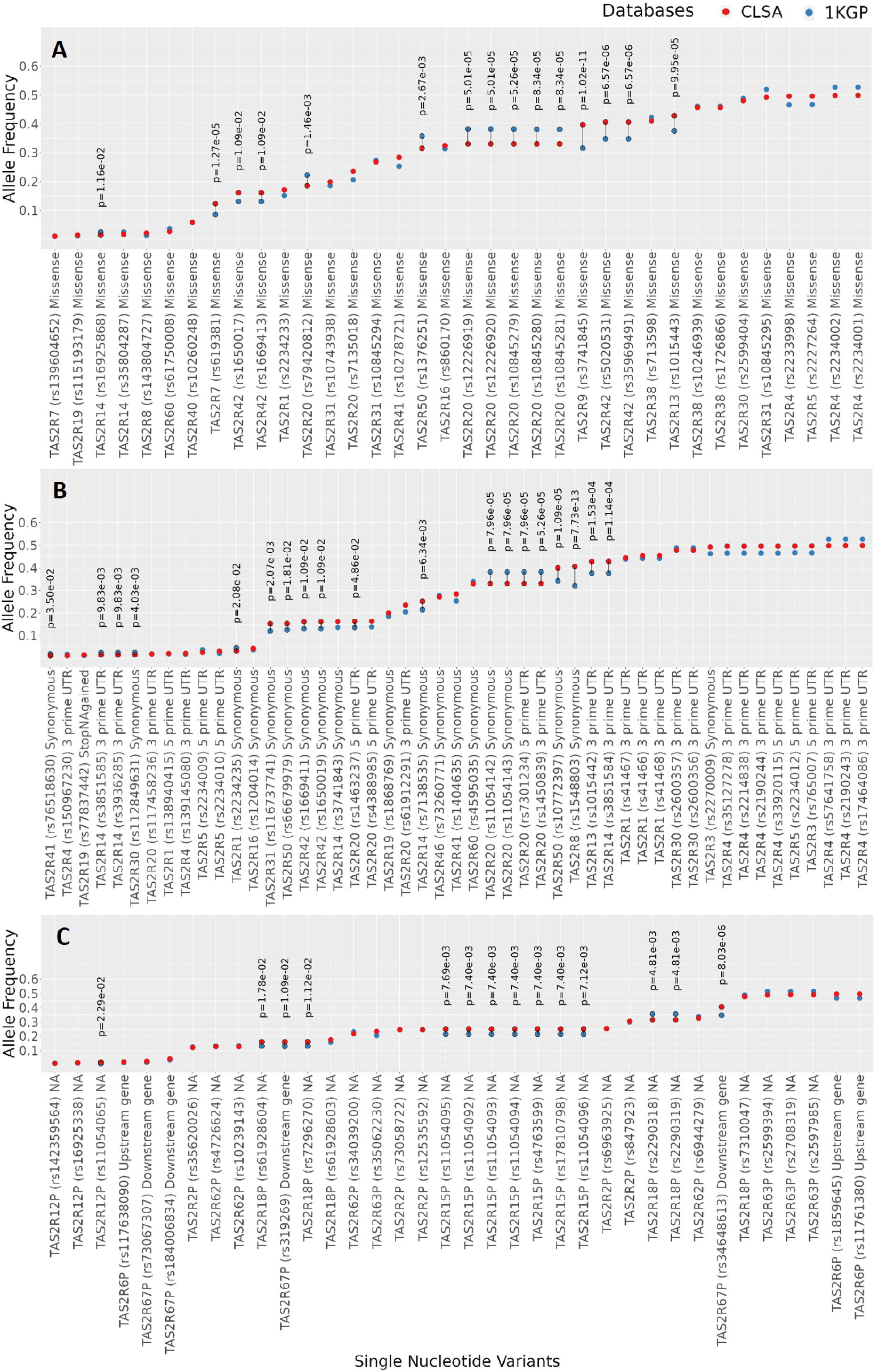
Allele distributions of *TAS2R* SNPs in CLSA and 1KGP. Dots represent the minor allele frequencies of each SNP within the European population: (**A**) Missense variants. (**B**) Non-missense variants. (**C**) Variants in *TAS2R* pseudogenes. Significant differences (adjusted p-value < 0.05, chi-squared test) are marked with a black line. NA: unknown functional consequence. See also Tables S1 and S2.

### Prevalence of oral health symptoms in the cohort

Following quality control of the cohort with available oral health data, 21,991 individuals were included, with a mean age of 63 years. The cohort consisted of 10,920 males and 11,071 females, as summarized in Table 1. Among the participants, 8.5% identified as current smokers, 44.3% as former smokers, and 47.1% as never having smoked. Table 2 provides a summary of the prevalence of oral health symptoms in the study cohort. Nearly 42.6% of the population reported no oral health problems. The most prevalent condition in the cohort was dry mouth, affecting 17.7% of individuals, followed by periodontal disease (14.6%) and decayed tooth (14.3%). Overall, 94.2% of the cohort reported their general oral health as excellent, very good, or good, while 5.8% rated it as fair or poor.

**Table 1.** Study population characteristics.

| Characteristic | Males (N = 10,920)<sup>a</sup> | Females (N = 11,071)<sup>a</sup> | P-value<sup>b</sup> |
| --- | --- | --- | --- |
| Age | 63 (55, 71) | 62 (55, 70) | 0.001 |
| Smoking |  |  | <0.001 |
| Never smoker | 4,816 (44%) | 5,553 (50%) |  |
| Former smoker | 5,089 (47%) | 4,659 (42%) |  |
| Current smoker | 1,013 (9.3%) | 859 (7.8%) |  |
| Self-reported general health |  |  | <0.001 |
| Excellent | 2,228 (20%) | 2,377 (21%) |  |
| Very good | 4,528 (42%) | 4,836 (44%) |  |
| Good | 3,235 (30%) | 2,976 (27%) |  |
| Fair | 763 (7.0%) | 726 (6.6%) |  |
| Poor | 155 (1.4%) | 149 (1.3%) |  |
| Education |  |  | <0.001 |
| Less than secondary school | 507 (4.7%) | 610 (5.5%) |  |
| Secondary school graduation | 908 (8.3%) | 1,177 (11%) |  |
| Some post-secondary education | 793 (7.3%) | 852 (7.7%) |  |
| Post-secondary degree/diploma | 8,689 (80%) | 8,419 (76%) |  |
| Household income |  |  | <0.001 |
| Less than \$20,000 | 335 (3.2%) | 638 (6.2%) | |
| \$20,000 to less than \$50,000 | 1,761 (17%) | 2,657 (26%) | |
| \$50,000 to less than \$100,000 | 3,779 (36%) | 3,618 (35%) | |
| \$100,000 to less than \$150,000 | 2,356 (23%) | 1,826 (18%) | |
| \$150,000 or more | 2,185 (21%) | 1,502 (15%) | |
| Brushing frequency |  |  | <0.001 |
| Less than once daily | 1,043 (9.6%) | 843 (7.6%) |  |
| Once daily | 2,802 (26%) | 1,472 (13%) |  |
| Twice daily or more | 7,059 (65%) | 8,749 (79%) |  |

<sup>a</sup> Median (IQR); n (%)
<sup>b</sup> Wilcoxon rank sum test; Pearson's Chi-squared test

**Table 2.** Prevalence of oral health symptoms in the CLSA European ancestry cohort.

| Characteristic | Males (N = 10,920) <sup>a</sup> | Females (N = 11,071) <sup>a</sup> | P-value <sup>b</sup> |
| --- | --- | --- | --- |
| General oral health |  |  | **<0.0001** |
| Excellent | 3,414 (31%) | 3,958 (36%) |  |
| Very good | 4,407 (40%) | 4,318 (39%) |  |
| Good | 2,399 (22%) | 2,210 (20%) |  |
| Fair | 545 (5.0%) | 447 (4.0%) |  |
| Poor | 141 (1.3%) | 120 (1.1%) |  |
| Uncomfortable eating in last 12 months |  |  | **<0.0001** |
| Often | 193 (1.8%) | 296 (2.7%) |  |
| Sometimes | 717 (6.6%) | 931 (8.4%) |  |
| Rarely | 2,135 (20%) | 2,197 (20%) |  |
| Never | 7,857 (72%) | 7,631 (69%) |  |
| Avoided eating in last 12 months |  |  | **<0.0001** |
| Often | 143 (1.3%) | 213 (1.9%) |  |
| Sometimes | 399 (3.7%) | 634 (5.7%) |  |
| Rarely | 1,031 (9.4%) | 1,313 (12%) |  |
| Never | 9,330 (85%) | 8,893 (80%) |  |
| Edentulism | 629 (5.8%) | 686 (6.2%) | 0.1160 |
| Toothache | 1,410 (13%) | 1,306 (12%) | **0.0460** |
| Cannot chew adequately | 799 (7.3%) | 933 (8.4%) | **0.0013** |
| Swelling in mouth | 465 (4.3%) | 535 (4.8%) | **0.0269** |
| Dry mouth | 1,689 (15%) | 2,196 (20%) | **<0.0001** |
| Burning mouth | 102 (0.9%) | 207 (1.9%) | **<0.0001** |
| Jaw muscles sore | 396 (3.6%) | 788 (7.1%) | **<0.0001** |
| Jaw joints painful | 404 (3.7%) | 868 (7.8%) | **<0.0001** |
| Natural tooth decayed | 1,798 (16%) | 1,352 (12%) | **<0.0001** |
| Natural tooth loose | 510 (4.7%) | 522 (4.7%) | 0.7087 |
| Natural tooth broken | 1,278 (12%) | 1,124 (10%) | **0.0017** |
| Gums around natural teeth are sore | 754 (6.9%) | 894 (8.1%) | **0.0006** |
| Gums around natural teeth bleed | 1,094 (10%) | 1,315 (12%) | **<0.0001** |
| Denture-related sores | 203 (1.9%) | 262 (2.4%) | **0.0062** |
| Teeth or dentures dirty | 386 (3.5%) | 239 (2.2%) | **<0.0001** |
| Bad breath | 823 (7.5%) | 735 (6.6%) | **0.0258** |
| Periodontal disease (bleeding gums or loose tooth) | 1,508 (14%) | 1,700 (15%) | **0.0007** |
| No oral health problems | 4,754 (44%) | 4,618 (42%) | 0.1600 |

<sup>a</sup> N (%)<sup>b</sup> Pearson's Chi-squared test. Significant values are bold (p < 0.05).

Certain conditions were significantly more prevalent in females (p<0.05), including discomfort while eating due to mouth problems, swelling in the mouth, periodontal diseases, denture-related sores, dry mouth, burning mouth, painful jaw joints, and sore jaw muscles. Notably, the latter three conditions were nearly twice as prevalent in females compared to males. Conversely, some conditions were more common in males (p<0.05): decayed tooth, broken tooth, toothache, bad breath, and dirty teeth or dentures. The prevalence of the remaining symptoms, including edentulism, loose tooth, and the overall presence or absence of symptoms, did not significantly differ between the sexes (p>0.05).

### *TAS2R* SNPs are associated with sore jaw muscles

The results of the chi-squared test revealed a significant association between 15 variants across 6 genes and sore jaw muscles (Table S3). Specifically, 9 variants in *TAS2R20* were associated with reduced prevalence of sore jaw muscles, while 6 variants in five other *TAS2R* genes were linked to an increased risk of this symptom. No other variants showed significant associations with sore jaw muscles or other oral health symptoms after applying the Bonferroni correction.

Logistic regression analysis, controlling for sex, age, brushing frequency, smoking status, education, household income, and self-reported general health, confirmed a significant association between the 15 variants and sore jaw muscles, as summarized in Table 3 and visualized in **Figure 2**. The RNAsnp web server predicted that seven of the 15 SNPs may affect mRNA secondary structure (**Figure S2**). Other variants and symptoms showed no significant association after adjusting for P-values. Detailed results are presented in Table S4, and additional plots for other symptoms are provided in **Figure S3**.

**Table 3.**
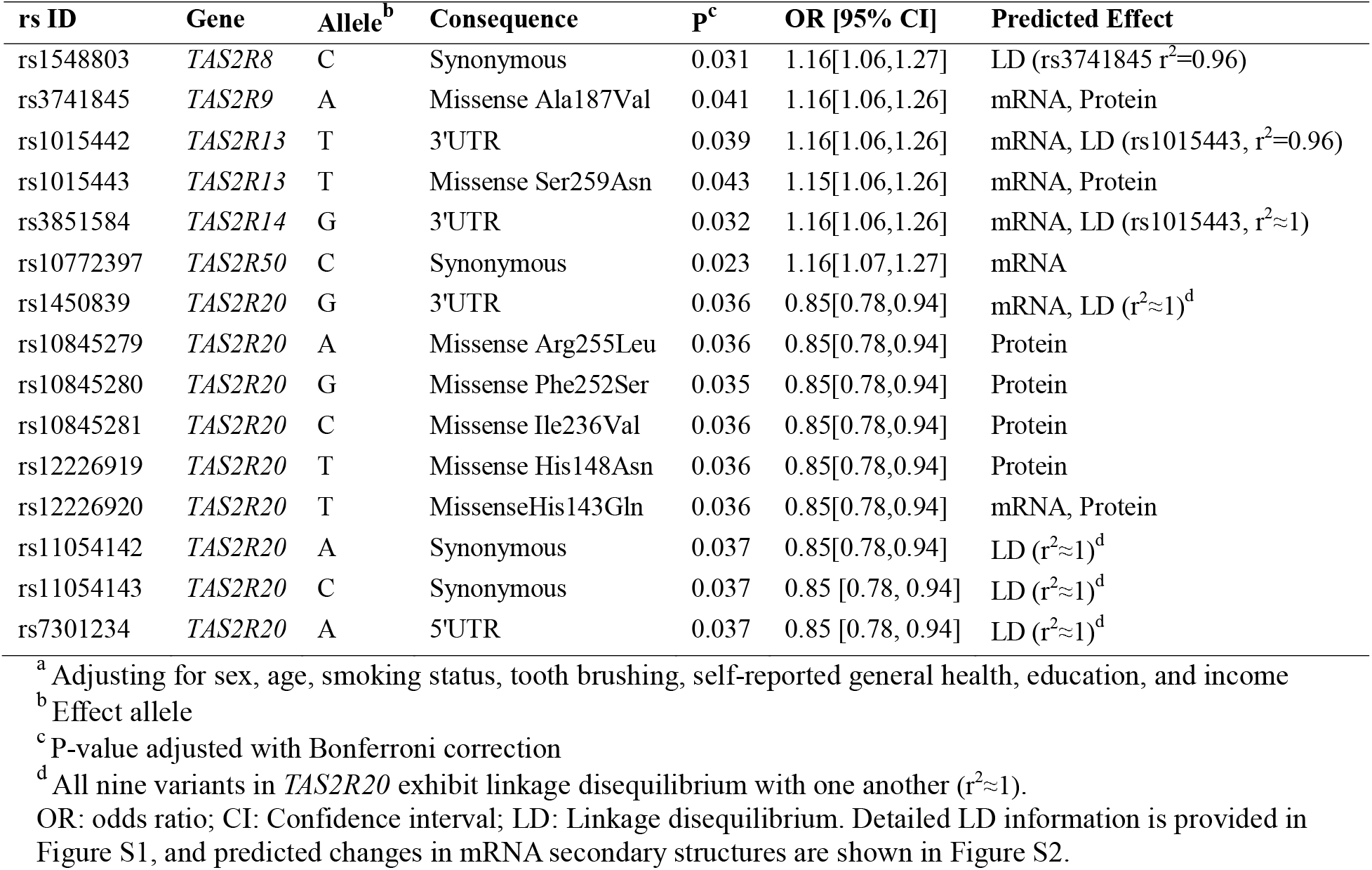
Significant associations between *TAS2R* variants and self-reported sore jaw muscles based on the logistic regression results.^a^.

**Figure 2.**
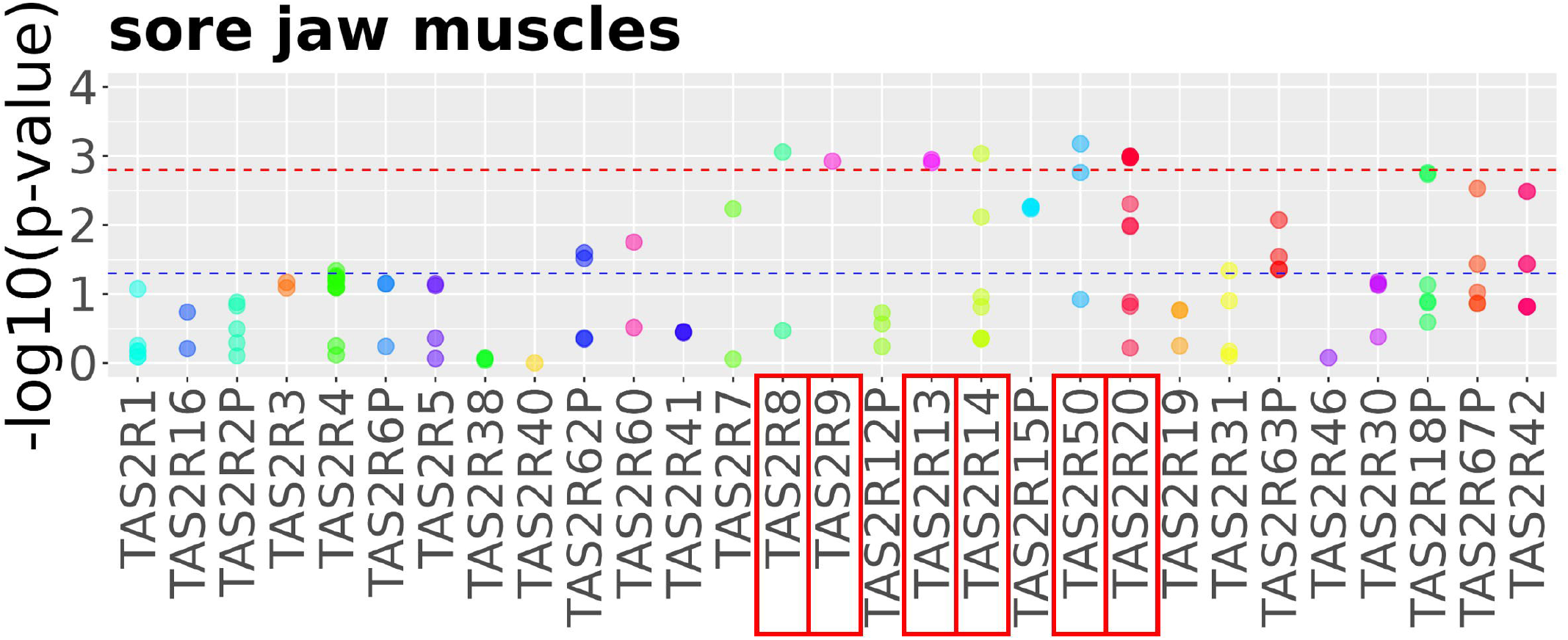
*TAS2R* SNPs are associated with self-reported sore jaw muscles. The blue line represents the significance threshold (p = 0.05), while the red line marks the Bonferroni-corrected threshold, with dots above this line indicating SNPs with significant associations (logistic regression). Genes containing these associated SNPs are highlighted with red rectangles. Additional Manhattan plots for other symptoms are available in **Figure S2**. For further details, refer to Table S4.

### Structure-function analysis of T2R20

Five of the seven missense SNPs associated with sore jaw muscles were in *TAS2R20*, prompting a focus on T2R20 for structure-function analysis. **Figure 3** presents 2D and 3D models of T2R20, highlighting the amino acids affected by the missense variants. The 3D model and ligand docking analysis revealed the locations of the CLSA variants within T2R20. Phenylalanine 252 was found in close proximity to the binding pocket (4.5 Å), potentially impacting ligand interactions, while the other variants were located approximately 10 Å from the binding site.

**Figure 3.**
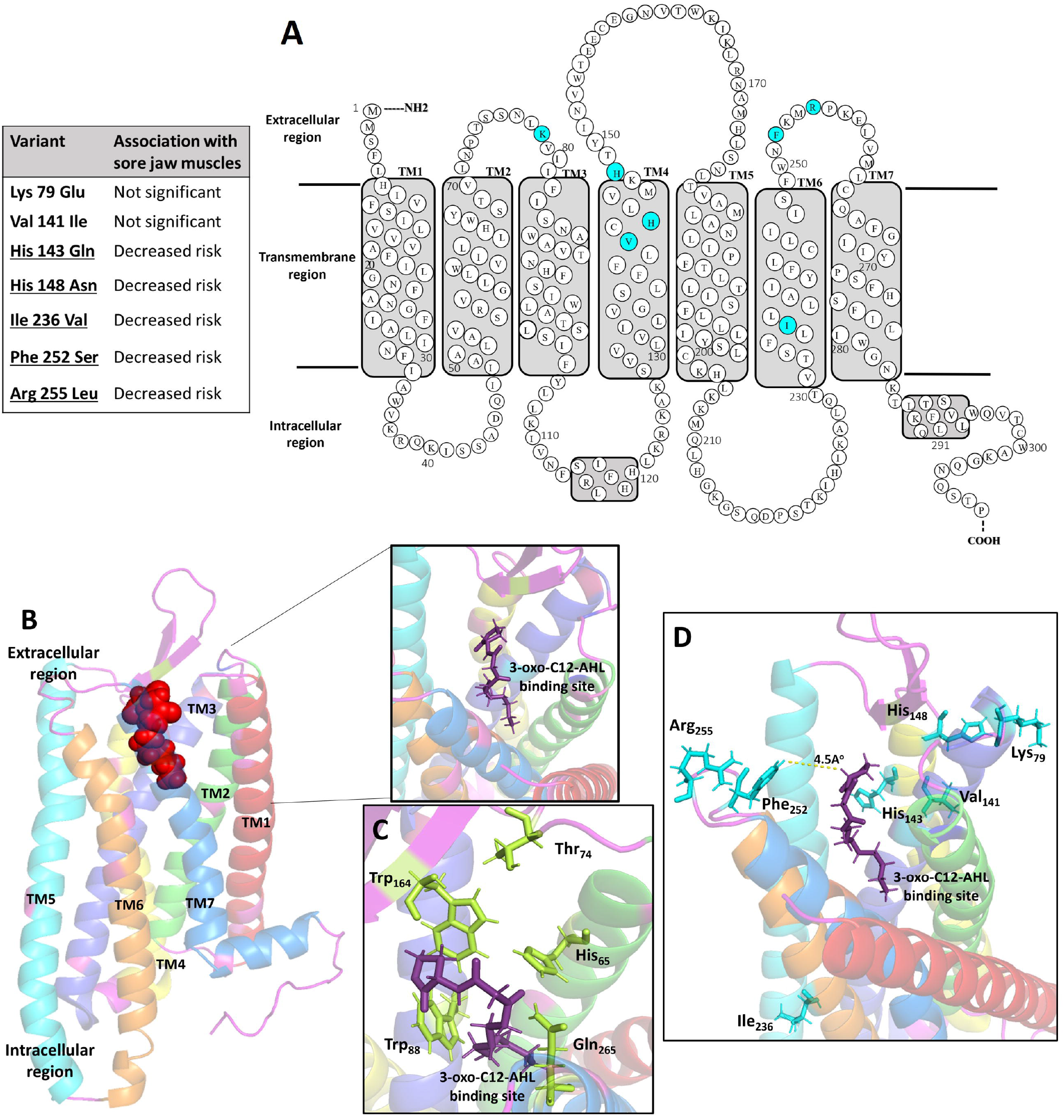
Structure-function analysis of T2R20. Models of T2R20 are adapted from previous studies.^15,17^ (**A**) 2D model of T2R20, highlighting amino acids affected by variants in the CLSA. Underlined variants are in linkage disequilibrium and associated with protection against sore jaw muscles. (**B-C**) 3D model of T2R20 from a prior publication, showing the binding site for bacterial AHL (red sphere-shaped structure), and the interacting amino acid residues.^15^ (**D**) T2R20 variants in the CLSA and their proximity to the binding site. Phenylalanine 252 is the closest residue to the binding site (4.5Å), with other variants within approximately 10 Å.

## DISCUSSION

This study investigated the allele distributions of 124 common *TAS2R* variants within the CLSA European ancestry cohort, identifying 48 SNPs with allele distributions differing from those in the 1KGP. Significant associations were identified between *TAS2R* variants and self-reported sore jaw muscles, particularly for *TAS2R20*. Structural analysis of T2R20 revealed two variants close to the binding site, suggesting that these genetic variants may influence the symptom by altering ligand interactions.

We observed extensive variation within *TAS2R* genes, with *TAS2R20* exhibiting the highest number of variants and *TAS2R15P* showing the greatest count of variants among the pseudogenes. Our results align with those of Wooding et al., who analyzed 1KGP data from approximately 2,500 individuals worldwide, excluding *TAS2R* pseudogenes.^34^ They reported the highest nucleotide diversity in *TAS2R20* and the lowest in *TAS2R39*. Additionally, our results corroborate a previous study on 911 Canadian young adults, which focused on rs713598 in *TAS2R38*—a tag SNP used to identify the supertaster genotype.^35^ They reported a frequency of the 49A allele at 55% among Caucasians, closely aligning with our finding of 59% among individuals of European ancestry.

The comparison of *TAS2R* genetic variants between the CLSA and 1KGP revealed generally similar allele distributions, with some notable differences, including in 11 missense variants. Six of these missense variants were in *TAS2R20*, all showing lower minor allele frequencies compared to the 1KGP. Previous studies have suggested a decrease in evolutionary pressure on human *TAS2Rs* during recent stages of evolution.^36-38^ Therefore, these differences between the cohorts could be attributed to factors such as population structure, founder effects, genetic bottlenecks, or genetic drift.^36,37,39^ Interestingly, five of the six *TAS2R20* alleles with lower frequencies in the CLSA were associated with a reduced risk of self-reported sore jaw muscles. This association suggests a potential functional relevance of these variants, which merits further investigation.

The findings of this study align with previous reports on oral health in the CLSA population.^40,41^ This study identified dry mouth as the most prevalent oral health issue among older Canadian adults, with a prevalence of 15% in men and 20% in women, followed by periodontal disease in 14% of men and 15% of women. Since early-stage periodontal disease is often asymptomatic, only severe cases could be identified through self-reported symptoms, defined as gingival bleeding or loose teeth.^42^ The 15% prevalence observed in this study aligns with a 2014 meta-analysis, which reported a 15-20% prevalence of severe periodontitis among North American adults aged 45 to 85.^43^

Sex differences in oral health symptoms were notable, with females experiencing a higher prevalence of xerostomia, burning mouth syndrome, denture-related sores, sore jaw muscles, and painful jaw joints. These differences may be attributed to underlying conditions such as medication use, hormonal changes, and autoimmune disorders such as Sjögren’s syndrome, which is more common in females.^44-48^ Decreased estrogen levels in perimenopausal women can lead to oral mucosal degeneration, increasing the risk of burning mouth syndrome.^49^ Sjögren’s syndrome, an autoimmune disorder causing oral cavity dryness, predominantly affects women.^46,47^ Xerostomia is also linked to *Candida* species, which can predispose individuals to denture-induced stomatitis.^50^ This is consistent with the higher prevalence of denture-related sores among females. Furthermore, sore jaw muscles and painful jaw joints were twice as prevalent in females, aligning with previous studies that report a greater prevalence of temporomandibular joint disorder (TMD) among females.^51-53^ These differences may partly be due to sex hormones, which can influence pain sensitivity.^44,45^ Additionally, TMD is associated with juvenile idiopathic arthritis, an autoimmune disease that has a greater incidence in female compared to male children and can lead to micrognathia and TMD in later adulthood. TMD is also linked to rheumatoid arthritis, an autoimmune disease that is two to three times more common in women.^48,54-57^

Our analysis of 124 SNPs identified significant associations between sore jaw muscles, the most prevalent manifestation of TMD,^58^ and 15 SNPs across six genes. This aligns with genome-wide association study (GWAS) data from the UK Biobank, which showed that six of these alleles, spanning five genes, were significantly more frequent in individuals with rheumatoid arthritis.^59^ The remaining variants were all located in *TAS2R20* and in linkage disequilibrium (Table S6).

Among the 15 associated SNPs, we identified four synonymous and four UTR variants, consistent with growing evidence that synonymous variants are linked to various diseases and can have effects comparable to those of nonsynonymous variants.^60,61^ Synonymous variants may impact mRNA structure, potentially altering its stability and translational speed, thus regulating post-transcriptional gene expression.^62,63^ Indeed, four of the associated synonymous and UTR variants may impact mRNA structure (**Figure S3)**. The remaining were in linkage disequilibrium with the associated missense variants (r^2^ > 0.9), suggesting that their phenotypic effects may be due to the polypeptide changes caused by the missense variants.

Among the five missense variants in *TAS2R20* linked to protection against sore jaw muscles, three were located in extracellular loops, where specific residues directly interact with ligands such as antibiotics and bacterial quorum-sensing molecules.^15,17^ 3D models revealed that Phe252, a residue in the third extracellular loop, is near the binding site and can potentially affect ligand interactions. Phe252 is in linkage disequilibrium with four other variants, meaning they are inherited together, which might significantly affect protein structure. The His148Asn variant is located in the second extracellular loop, at the interface with the transmembrane helix. This substitution might make the local structure more rigid due to Asn’s smaller and less flexible side chain compared to His, potentially affecting the movement and positioning of the extracellular loop and ligand interactions.

TMD encompasses a range of neuromuscular and musculoskeletal disorders impacting the temporomandibular joint, masticatory muscles, and/or related tissues, resulting in symptoms such as pain, difficulty chewing, and, in severe cases, restricted jaw movement.^58,64,65^ It is influenced by multiple risk factors, including malocclusion, oral parafunctions such as bruxism, psychological factors like depression and anxiety, and iatrogenic injuries.^53^ Elevated inflammatory markers, including interleukins and TNF-α, have been detected in the blood, masseter muscle extracellular fluid, and saliva of individuals with TMD.^52,66^ Additionally, specific bacterial species, particularly *Staphylococcus aureus*, have been implicated in TMD.^67,68^

The SNPs associated with sore jaw muscles were located in *TAS2R8, 9, 13, 14, 20*, and *50*, potentially due to the specific roles and expression patterns of these receptors. *TAS2R13, 14, 20*, and *50* are expressed at moderate to high levels in extraoral tissues.^12,69^ Notably, T2R20 and T2R14 detect bacterial quorum-sensing molecules and mediate immune response.^17,70^ A previous study showed that T2R14 responds to molecules from *S. aureus* by inducing inflammatory markers and inhibiting bacterial growth.^71^ Thus, *TAS2R*s may contribute to TMD pathogenesis by modulating the immune response to pathogens and triggering the release of inflammatory biomarkers such as TNF-α and interleukins.^15-17,70,71^

Our study found no significant association with other self-reported oral health symptoms. Research on this subject is limited and primarily focuses on dental caries and *TAS2R38* variants, which have yielded contradictory results.^18,28-31^ A recent meta-analysis of genome-wide association studies also found no significant association between *TAS2R* SNPs and early childhood caries.^24^ However, their transcriptome-wide analysis identified a significant association with the expression of three *TAS2R*s, including *T2R14*, which was associated with sore jaw muscles in our study. In terms of periodontal disease, Khimsuksri et al. found a lower risk of periodontitis (pocket depth ≥ 5 mm) in individuals with the *TAS2R38* AVI non-taster genotype.^28^ Conversely, Kaur et al. observed no significant association between *TAS2R* variants and the Periodontal Screening and Recording (PSR) index, which evaluates gingival bleeding and pocket depth, consistent with our findings.^72^ We defined periodontal disease as the presence of gingival bleeding and/or loose teeth, supporting the comparable outcomes.

Findings of this study enhance our understanding of genetic risk factors for oral health and highlight the potential role of *TAS2R*s in the pathogenesis of TMD. These receptors, especially the disease-associated residues, could be promising targets for the development of new pharmacological agents. Moreover, identifying genetic variants linked to these conditions could aid in recognizing patients at higher risk, enabling more personalized medicine and tailored treatment plans.

## RESOURCE AVAILABILITY

### Lead contact

Further information should be directed to Dr. Carol A. Hitchon.

### Materials availability

This study did not generate new unique reagents.

### Data and code availability

- Data: This study used the CLSA genome-wide genetic data release (version 3) and Baseline Questionnaire data. Data are available from the Canadian Longitudinal Study on Aging (www.clsa-elcv.ca) for researchers who meet the criteria for access to de-identified CLSA data. Allele frequencies from the 1KGP were obtained through the Ensembl Variant Effect Predictor (VEP). The DOI is provided in the key resources table.
- Code: The source code of this article has been deposited at https://github.com/mshafizadeh/TAS2R_OralHealth and is publicly available as of the date of publication.
- Any additional information will be made available upon request from the lead contact.

### Limitations of the Study

There are several limitations to the current study. Although the comparison with 1KGP was conducted due to its availability, the small sample sizes of subpopulations limited our ability to account for potential outliers. While we identified some differences between the datasets, exploring their evolutionary origins was not within the scope of this study. Additionally, our analyses focused on individuals of European ancestry, limiting the extension of the findings to other ethnic groups. The self-reported data may introduce potential bias, as symptoms could not be validated through clinical exams due to the large sample size. Nevertheless, these oral health indicators have demonstrated adequate validity and reliability, and the large sample size enhances the study’s statistical power.^73,74^ Another limitation is the cross-sectional design, which prevents determining causality between the variants and the symptom. This limitation could be addressed in future longitudinal studies as more CLSA follow-up data become available. Furthermore, the number of SNPs detected in different genes varied, which may contribute to the differing number of associated SNPs. While the study identified significant associations, it did not account for potential interactions with environmental factors or other genetic variants. Finally, although the structure-function analysis is suggestive, further experimental validation is required to confirm the underlying biological mechanisms.

## ACKNOWLEDGMENTS

This study is funded by a Canadian Institutes of Health Research (CIHR) Catalyst Grant (ACD 187255) and an NSERC (Natural Sciences and Engineering Research Council of Canada) Discovery Grant (RGPIN-2020-05670). We thank M. Wasif Khan for advice on statistical analysis.

This research was made possible using the data/biospecimens collected by the Canadian Longitudinal Study on Aging (CLSA). Funding for the Canadian Longitudinal Study on Aging (CLSA) is provided by the Government of Canada through the Canadian Institutes of Health Research (CIHR) under grant reference: LSA 94473 and the Canada Foundation for Innovation, as well as the following provinces, Newfoundland, Nova Scotia, Quebec, Ontario, Manitoba, Alberta, and British Columbia. This research has been conducted using the CLSA Genome-wide Genetic Data Release (version 3) and Comprehensive Baseline Dataset (version 5), under the Application ID 2006008. The CLSA is led by Drs. Parminder Raina, Christina Wolfson and Susan Kirkland.

## Disclaimer

The opinions expressed in this manuscript are the author’s own and do not reflect the views of the Canadian Longitudinal Study on Aging.

## AUTHOR CONTRIBUTIONS

Conceptualization, C.A.H., P.C.; Methodology, M.S., S.A., B.D., P.S.J., C.S.; Investigation, M.S., V.B.; Writing – Original Draft, M.S.; Writing – Review and Editing, M.S., V.B., R.P.B., P.C., C.A.H., B.D., C.S., P.S.J., S.A.; Visualization, M.S., V.B.; Funding Acquisition, C.A.H., P.C.; Supervision, C.A.H., P.C., R.P.B. All authors have read and agreed to the published version of the manuscript.

## DECLARATION OF INTERESTS

The authors declare that they have no competing interests related to this work.

## FIGURE TITLES AND LEGENDS

## STAR⍰METHODS

### KEY RESOURCES TABLE

STAR<sup>®</sup> METHODS KEY RESOURCES TABLE
| Resource | Source | Identifier |
| --- | --- | --- |
| PLINK version v.1.9 | Purcell et al.<sup>75</sup> | https://zzz.bwh.harvard.edu/plink/data.shtml#plink |
| R version 4.3.2 | R Core Team<sup>76</sup> | https://www.r-project.org/ |
| NCBI databases | Sayers et al.<sup>77</sup> | https://www.ncbi.nlm.nih.gov/ |
| Ensembl Variant Effect Predictor (VEP) | McLaren et al.<sup>78</sup> | https://www.ensembl.org/info/docs/tools/vep/index.html |
| UK Biobank Imputed Version 3 | Neale Lab<sup>59</sup> | http://www.nealelab.is/uk-biobank |
| I-TASSER | Yang et al.<sup>79</sup> | https://zhanggroup.org/I-TASSER/ |

### Experimental MODEL AND SUBJECT DETAILS

Participant details can be found in Table 1. The inclusion and exclusion criteria used are provided in the sections below.

## METHOD DETAILS

### Study population and data preprocessing

The study design and statistical analysis process are summarized in **Figure 4**. Our team received ethics approval from the University of Manitoba’s Health Research Ethics Board (HREB HS26021-H2023:173) and obtained access to the CLSA questionnaire data and genome-wide genetic dataset version 3 (CLSA Application ID: 2006008). Details of the CLSA protocol, including inclusion/exclusion criteria, sampling strategy, data collection, and procedures, are available for download under the Researchers section of the CLSA website (www.clsa-elcv.ca). In brief, individuals aged 45-85 at baseline were recruited from multiple sites across Canada and followed with standardized data collection, including demographics and lifestyle/behavioral measures. A subset consented to provide biosamples, including blood for genotyping. Baseline and genetic data were used for this analysis. All CLSA participants consented to the use of their data, which is securely stored on our lab server. The genetic dataset includes information from 26,622 participants, who were genotyped for 794,409 variants using the Affymetrix Axiom array, along with an additional 308 million imputed variants from the TOPMed reference panel.^32^ The positions of *TAS2R* genes in the GRCh38 genome assembly were obtained from the National Center for Biotechnology Information (NCBI) Gene database.^77^ We used PLINK (v1.9)^75^ to extract SNPs in *TAS2R* genes based on the GRCh38 positions. Variant annotations were obtained from the Ensembl Variant Effect Predictor (VEP).^78^

**Figure 4.**
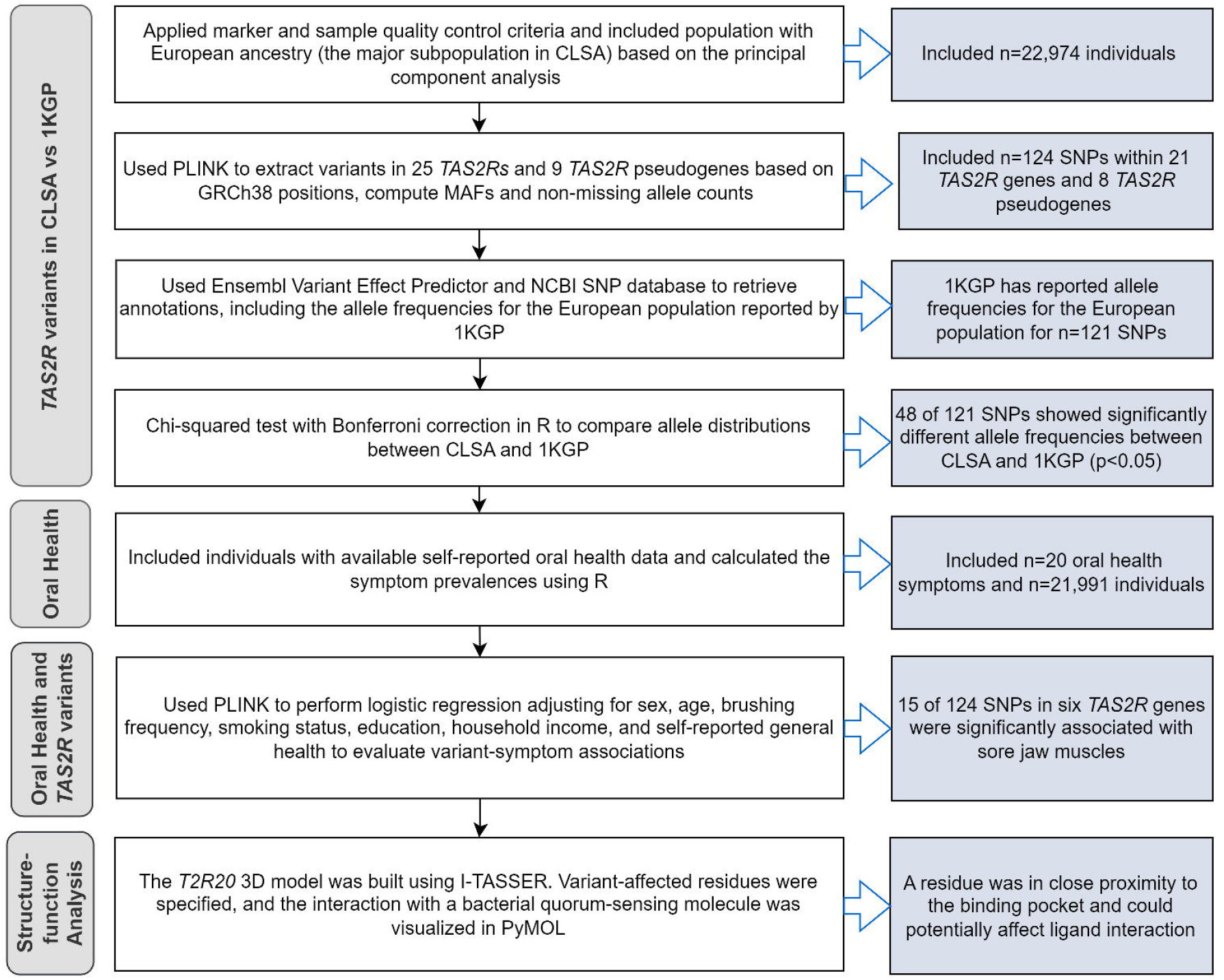
Study pipeline for analysis of *TAS2R* genetic variants in the CLSA cohort. SNP: single nucleotide polymorphism; MAF: minor allele frequency; 1KGP: 1000 Genomes Project.

### Eligibility criteria

Sample and marker quality control was applied using PLINK according to protocols established in prior CLSA publications.^32^ Genetic markers were excluded based on the following criteria: genotype frequency discordance between batches and across control replicates; deviation from Hardy-Weinberg equilibrium (HWE) with a p-value threshold of 3.15 × 10^−10^; insertions/deletions (indels); genotype missingness greater than 5%; and SNPs with minor allele frequency (MAF) ≤ 0.01. Sample exclusions were based on the following: individuals with familial relatedness of 3^rd^ degree or closer (determined by identity-by-descent (IBD) analysis); heterozygosity outliers; discordance between self-reported and chromosomal sex; and samples with more than 5% missing genotypes. Furthermore, imputed variants with a quality score above 0.8 were included in the analysis.^80^ To minimize the impact of population stratification in this study, we included only individuals of European ancestry, selected as previously described by the CLSA.^32^

### Oral health data in CLSA

Oral health data were obtained from the CLSA self-reported questionnaire, collected between 2011 and 2015. The CLSA oral health questionnaire, derived from the Canadian Community Health Survey 2.1, is based on self-reported oral health indicators with demonstrated good construct and concurrent validity, internal consistency, and reliability greater than 0.7.^73,81^ The questions and response values are summarized in Table S5. In addition to the 20 variables directly obtained from the questionnaire, periodontal disease was defined as the report of at least one loose tooth and/or bleeding gingiva. This definition was validated against prevalence data from the literature. For the genetic association analyses, using PLINK, categorical variables with more than two categories were dichotomized as follows: general oral health (good: excellent, very good, good vs. poor: fair, poor); and avoided eating and uncomfortable eating (yes: often, sometimes vs. no: rarely, never).

## STATISTICAL ANALYSIS

### *TAS2R* allele distributions in CLSA and 1KGP

Allele frequencies and non-missing allele counts were calculated using PLINK for the CLSA population of European ancestry. To validate the results, chi-squared tests were applied using R (v4.3.2)^76^ to compare the allelic distributions with those reported for the European population in the 1000 Genomes Project (1KGP).^82^ Bonferroni correction was applied by multiplying p-values by the number of independent markers in linkage equilibrium (r^2^ < 0.1 within a 1000 kb window, n=35).^32^ P<0.05 was considered statistically significant. Scatter plots of allele frequencies were generated using the ggplot2 package in R.^83^ Pairwise linkage disequilibrium between the SNPs was visualized using the corrplot package.

### *TAS2R* SNPs and oral health symptoms

Individuals with available oral health data were included in the genetic association analysis, with 87 excluded due to conflicting data, as they reported both an absence of natural tooth and tooth-related problems simultaneously. The prevalence of oral health symptoms was summarized usingR. Patients with edentulism were excluded from the analysis of tooth-related phenotypes. Chi-squared tests were performed using PLINK to compare allele distributions between cases and controls. Logistic regression analysis was conducted using an additive genetic model in PLINK to identify variant-disease associations, adjusting for sex, age, smoking status, brushing frequency, self-reported general health, education, and household income. Correlation analysis was performed to ensure the absence of multicollinearity between the covariates. Manhattan plots were generated using ggplot2 package to visualize the results. P-values were adjusted using Bonferroni correction based on the number of independent markers multiplied by the number of independent phenotypes, with a significance threshold of P<0.05. Given the moderate to strong correlations observed among most phenotypes (Phi coefficient > 0.1, **Figure S4**), the number of independent phenotypes was considered as 1 for the Bonferroni correction.^84^ RNAsnp web server was used to predict the impact of the variants on mRNA structure.^85^ To validate the findings in an independent cohort, the associated variants were examined in genome-wide association study (GWAS) results from the UK Biobank Imputed Version 3 dataset, generated by the Neale Lab, to determine whether they are linked to relevant phenotypes (Table S6).^59^

### Structure-function analysis

To illustrate the impact of the variants on protein structure, we used a 2-dimensional model of T2R20 from a previous publication,^17^ highlighting the amino acids affected by the variants.

Additionally, we employed a 3-dimensional model of T2R20, constructed using I-TASSER as described in a previous study.^15^ I-TASSER is a hierarchical method for predicting protein structure.^79^ Energy minimization of the T2R20 structure and ligand docking with the bacterial quorum sensing molecule 3-oxo-C12-Acyl homoserine lactone (AHL) were performed using Maestro v11.0 (Schrödinger, LLC, New York), as previously described.^15^ The interactions between T2R20 residues and AHL within the binding pocket, along with residues affected by the CLSA variants, were visualized using PyMOL molecular graphics software v3.0 (Schrödinger, LLC, New York). The distances of residues from the binding site were calculated using the measurement wizard in PyMOL.

## SUPPLEMENTAL INFORMATION

**Document S1**. Tables S1, S2, S5, and S6, and Figures S1–4.

**Table S3**. Excel file containing the chi-squared test results comparing allele distributions of *TAS2R*

SNPs in individuals with and without self-reported oral health symptoms, related to Figure 2. **Table S4**. Excel file containing the logistic regression results on the association between *TAS2R* SNPs and self-reported oral health symptoms in the CLSA, adjusted for confounding variables, related to Figure 2 and Table 3.

## Notes

### Competing Interest Statement

The authors have declared no competing interest.

## REFERENCES

1. Braschi, B., Denny, P., Gray, K., Jones, T., Seal, R., Tweedie, S., Yates, B., and Bruford, E. (2019). Genenames.org: the HGNC and VGNC resources in 2019. Nucleic Acids Research 47, D786–D792. 10.1093/nar/gky930.

2. Behrens, M., Foerster, S., Staehler, F., Raguse, J.D., and Meyerhof, W. (2007). Gustatory expression pattern of the human TAS2R bitter receptor gene family reveals a heterogenous population of bitter responsive taste receptor cells. The Journal of neuroscience : the official journal of the Society for Neuroscience 27, 12630–12640. 10.1523/jneurosci.1168-07.2007.

3. Meyerhof, W., Batram, C., Kuhn, C., Brockhoff, A., Chudoba, E., Bufe, B., Appendino, G., and Behrens, M. (2010). The molecular receptive ranges of human TAS2R bitter taste receptors. Chemical senses 35, 157–170. 10.1093/chemse/bjp092.

4. Hoon, M.A., Adler, E., Lindemeier, J., Battey, J.F., Ryba, N.J., and Zuker, C.S. (1999). Putative mammalian taste receptors: a class of taste-specific GPCRs with distinct topographic selectivity. Cell 96, 541–551. 10.1016/s0092-8674(00)80658-3.

5. Conte, C., Ebeling, M., Marcuz, A., Nef, P., and Andres-Barquin, P.J. (2002). Identification and characterization of human taste receptor genes belonging to the TAS2R family. Cytogenetic and genome research 98, 45–53. 10.1159/000068546.

6. Seal RL B.B.,, Gray K, Jones TEM, Tweedie S, Haim-Vilmovsky L, Bruford EA. Genenames.org: the HGNC resources in 2023. Nucleic Acids Res. PMID: 36243972 DOI: 10.1093/nar/gkac888.

7. Balakirev, E.S., and Ayala, F.J. (2003). Pseudogenes: are they “junk” or functional DNA? Annual review of genetics 37, 123–151. 10.1146/annurev.genet.37.040103.103949.

8. Tam, O.H., Aravin, A.A., Stein, P., Girard, A., Murchison, E.P., Cheloufi, S., Hodges, E., Anger, M., Sachidanandam, R., Schultz, R.M., and Hannon, G.J. (2008). Pseudogene-derived small interfering RNAs regulate gene expression in mouse oocytes. Nature 453, 534–538. 10.1038/nature06904.

9. Ma, Y., Chen, Z., and Yu, J. (2021). Pseudogenes and their potential functions in hematopoiesis. Experimental hematology 103, 24–29. 10.1016/j.exphem.2021.09.001.

10. Gil, S., Coldwell, S., Drury, J.L., Arroyo, F., Phi, T., Saadat, S., Kwong, D., and Chung, W.O. (2015). Genotype-specific regulation of oral innate immunity by T2R38 taste receptor. Molecular Immunology 68, 663–670. 10.1016/j.molimm.2015.10.012.

11. Kim, U., Wooding, S., Ricci, D., Jorde, L.B., and Drayna, D. (2005). Worldwide haplotype diversity and coding sequence variation at human bitter taste receptor loci. Human mutation 26, 199–204. 10.1002/humu.20203.

12. Jaggupilli, A., Singh, N., Upadhyaya, J., Sikarwar, A.S., Arakawa, M., Dakshinamurti, S., Bhullar, R.P., Duan, K., and Chelikani, P. (2017). Analysis of the expression of human bitter taste receptors in extraoral tissues. Molecular and cellular biochemistry 426, 137–147. 10.1007/s11010-016-2902-z.

13. Maurer, S., Wabnitz, G.H., Kahle, N.A., Stegmaier, S., Prior, B., Giese, T., Gaida, M.M., Samstag, Y., and Hänsch, G.M. (2015). Tasting Pseudomonas aeruginosa Biofilms: Human Neutrophils Express the Bitter Receptor T2R38 as Sensor for the Quorum Sensing Molecule N-(3-Oxododecanoyl)-l-Homoserine Lactone. Frontiers in immunology 6, 369. 10.3389/fimmu.2015.00369.

14. Tran, H.T.T., Herz, C., Ruf, P., Stetter, R., and Lamy, E. (2018). Human T2R38 Bitter Taste Receptor Expression in Resting and Activated Lymphocytes. Frontiers in immunology 9, 2949. 10.3389/fimmu.2018.02949.

15. Jaggupilli, A., Singh, N., De Jesus, V.C., Gounni, M.S., Dhanaraj, P., and Chelikani, P. (2019). Chemosensory bitter taste receptors (T2Rs) are activated by multiple antibiotics. The FASEB Journal 33, 501–517. 10.1096/fj.201800521RR.

16. Jaggupilli, A., Howard, R., Aluko, R.E., and Chelikani, P. (2019). Advanced glycation end-products can activate or block bitter taste receptors. Nutrients 11, 1317. 10.3390/nu11061317.

17. Jaggupilli, A., Singh, N., Jesus, V.C.D., Duan, K., and Chelikani, P. (2018). Characterization of the binding sites for bacterial acyl homoserine lactones (AHLs) on human bitter taste receptors (T2Rs). ACS Infectious Diseases 4, 1146–1156. 10.1021/acsinfecdis.8b00094.

18. de Jesus, V.C., Mittermuller, B.-A., Hu, P., Schroth, R.J., and Chelikani, P. (2022). Genetic variants in taste genes play a role in oral microbial composition and severe early childhood caries. Iscience 25. 10.1016/j.isci.2022.105489.

19. de Jesus, V.C., Singh, M., Schroth, R.J., Chelikani, P., and Hitchon, C.A. (2021). Association of Bitter Taste Receptor T2R38 Polymorphisms, Oral Microbiota, and Rheumatoid Arthritis. Current issues in molecular biology 43, 1460–1472. 10.3390/cimb43030103.

20. Dotson, C.D., Shaw, H.L., Mitchell, B.D., Munger, S.D., and Steinle, N.I. (2010). Variation in the gene TAS2R38 is associated with the eating behavior disinhibition in Old Order Amish women. Appetite 54, 93–99. 10.1016/j.appet.2009.09.011.

21. Eriksson, L., Esberg, A., Haworth, S., Holgerson, P.L., and Johansson, I. (2019). Allelic Variation in Taste Genes Is Associated with Taste and Diet Preferences and Dental Caries. Nutrients 11. 10.3390/nu11071491.

22. Yang, X., Wang, J., Hong, H., Feng, X., Zhang, X., and Song, J. (2024). The association between diets and periodontitis: a bidirectional two-sample Mendelian randomization study. Front Genet 15. 10.3389/fgene.2024.1398101.

23. Petrenya, N., Brustad, M., Hopstok, L.A., Holde, G.E., and Jönsson, B. (2024). Empirically derived dietary patterns in relation to periodontitis and number of teeth among Norwegian adults. Public Health Nutrition 27, e27. e27. 10.1017/S1368980023002690.

24. Orlova, E., Dudding, T., Chernus, J.M., Alotaibi, R.N., Haworth, S., Crout, R.J., Lee, M.K., Mukhopadhyay, N., Feingold, E., Levy, S.M., et al. (2023). Association of Early Childhood Caries with Bitter Taste Receptors: A Meta-Analysis of Genome-Wide Association Studies and Transcriptome-Wide Association Study. Genes 14, 59. 10.3390/genes14010059.

25. Keller, M., Liu, X., Wohland, T., Rohde, K., Gast, M.-T., Stumvoll, M., Kovacs, P., Tönjes, A., and Böttcher, Y. (2013). TAS2R38 and Its Influence on Smoking Behavior and Glucose Homeostasis in the German Sorbs. PLOS ONE 8, e80512. 10.1371/journal.pone.0080512.

26. Mao, Z., Cheng, W., Li, Z., Yao, M., and Sun, K. (2023). Clinical Associations of Bitter Taste Perception and Bitter Taste Receptor Variants and the Potential for Personalized Healthcare. Pharmacogenomics and personalized medicine 16, 121–132. 10.2147/pgpm.s390201.

27. Pitiphat, W., Merchant, A.T., Rimm, E.B., and Joshipura, K.J. (2003). Alcohol Consumption Increases Periodontitis Risk. Journal of dental research 82, 509–513. 10.1177/154405910308200704.

28. Khimsuksri, S., Paphangkorakit, J., Pitiphat, W., and Coldwell, S.E. (2022). TAS2R38 polymorphisms and oral diseases in Thais: a cross-sectional study. BMC oral health 22, 21. 10.1186/s12903-022-02043-2.

29. Kiliç, M., Gurbuz, T., Kahraman, C.Y., Cayir, A., Bilgiç, A., and Kurt, Y. (2022). Relationship between the TAS2R38 and TAS1R2 polymorphisms and the dental status in obese children. Dental and medical problems 59, 233–240. 10.17219/dmp/143252.

30. Wendell, S., Wang, X., Brown, M., Cooper, M.E., DeSensi, R.S., Weyant, R.J., Crout, R., McNeil, D.W., and Marazita, M.L. (2010). Taste genes associated with dental caries. Journal of dental research 89, 1198–1202. 10.1177/0022034510381502.

31. AlMarshad, L.K., AlJobair, A.M., Al-Anazi, M.R., Bohol, M.F.F., Wyne, A.H., and Al-Qahtani, A.A. (2021). Association of polymorphisms in genes involved in enamel formation, taste preference and immune response with early childhood caries in Saudi pre-school children. Saudi journal of biological sciences 28, 2388–2395. 10.1016/j.sjbs.2021.01.036.

32. Forgetta, V., Li, R., Darmond-Zwaig, C., Belisle, A., Balion, C., Roshandel, D., Wolfson, C., Lettre, G., Pare, G., Paterson, A.D., et al. (2022). Cohort profile: genomic data for 26 622 individuals from the Canadian Longitudinal Study on Aging (CLSA). BMJ open 12, e059021. 10.1136/bmjopen-2021-059021.

33. Raina, P.S., Wolfson, C., Kirkland, S.A., Griffith, L.E., Oremus, M., Patterson, C., Tuokko, H., Penning, M., Balion, C.M., Hogan, D., et al. (2009). The Canadian Longitudinal Study on Aging (CLSA). Canadian Journal on Aging / La Revue canadienne du vieillissement 28, 221–229. 10.1017/S0714980809990055.

34. Wooding, S.P., and Ramirez, V.A. (2022). Global population genetics and diversity in the TAS2R bitter taste receptor family. Front Genet 13, 952299. 10.3389/fgene.2022.952299.

35. Khataan, N.H., Stewart, L., Brenner, D.M., Cornelis, M.C., and El-Sohemy, A. (2009). TAS2R38 genotypes and phenylthiocarbamide bitter taste perception in a population of young adults. Journal of nutrigenetics and nutrigenomics 2, 251–256. 10.1159/000297217.

36. Kulichová, I., Mouterde, M., Mokhtar, M.G., Diallo, I., Tříska, P., Diallo, Y.M., Hofmanová, Z., Poloni, E.S., and Černý, V. (2022). Demographic history was a formative mechanism of the genetic structure for the taste receptor TAS2R16 in human populations inhabiting Africa’s Sahel/Savannah Belt. American journal of biological anthropology 177, 540–555. 10.1002/ajpa.24448.

37. Roudnitzky, N., Risso, D., Drayna, D., Behrens, M., Meyerhof, W., and Wooding, S.P. (2016). Copy Number Variation in TAS2R Bitter Taste Receptor Genes: Structure, Origin, and Population Genetics. Chemical senses 41, 649–659. 10.1093/chemse/bjw067.

38. Wooding, S.P., and Ramirez, V.A. (2022). Global population genetics and diversity in the TAS2R bitter taste receptor family. Front Genet 13. 10.3389/fgene.2022.952299.

39. Kim, U.K., and Drayna, D. (2005). Genetics of individual differences in bitter taste perception: lessons from the PTC gene. Clinical genetics 67, 275–280. 10.1111/j.1399-0004.2004.00361.x.

40. Bassim, C., Mayhew, A.J., Ma, J., Kanters, D., Verschoor, C.P., Griffith, L.E., and Raina, P. (2020). Oral Health, Diet, and Frailty at Baseline of the Canadian Longitudinal Study on Aging. Journal of the American Geriatrics Society 68, 959–966. 10.1111/jgs.16377.

41. De Rubeis, V., Jiang, Y., de Groh, M., Dufour, L., Bronsard, A., Morrison, H., Butt, F., and Walker Bassim, C. (2023). Oral Health Problems among Canadians Aged 45 to 85: Data from the Canadian Longitudinal Study on Aging Baseline Survey (2011-2015). International journal of environmental research and public health 20. 10.3390/ijerph20085533.

42. Gasner, N.S., and Schure, R.S. (2024). Periodontal disease. In StatPearls, (Treasure Island (FL): StatPearls Publishing). https://www.ncbi.nlm.nih.gov/books/NBK554590/.

43. Kassebaum, N.J., Bernabé, E., Dahiya, M., Bhandari, B., Murray, C.J., and Marcenes, W. (2014). Global burden of severe periodontitis in 1990-2010: a systematic review and meta-regression. Journal of dental research 93, 1045–1053. 10.1177/0022034514552491.

44. Fillingim, R.B., King, C.D., Ribeiro-Dasilva, M.C., Rahim-Williams, B., and Riley, J.L., 3rd (2009). Sex, gender, and pain: a review of recent clinical and experimental findings. The journal of pain 10, 447–485. 10.1016/j.jpain.2008.12.001.

45. Craft, R.M. (2007). Modulation of pain by estrogens. Pain 132 Suppl 1, S3–s12. 10.1016/j.pain.2007.09.028.

46. Qin, B., Wang, J., Yang, Z., Yang, M., Ma, N., Huang, F., and Zhong, R. (2015). Epidemiology of primary Sjögren’s syndrome: a systematic review and meta-analysis. Annals of the rheumatic diseases 74, 1983–1989. 10.1136/annrheumdis-2014-205375.

47. Retamozo, S., Acar-Denizli, N., Horváth, I.F., Ng, W.F., Rasmussen, A., Dong, X., Li, X., Baldini, C., Olsson, P., Priori, R., et al. (2021). Influence of the age at diagnosis in the disease expression of primary Sjögren syndrome. Analysis of 12,753 patients from the Sjögren Big Data Consortium. Clinical and experimental rheumatology 39 Suppl 133, 166–174. 10.55563/clinexprheumatol/egnd1i.

48. Gabriel, S.E. (2001). The epidemiology of rheumatoid arthritis. Rheumatic diseases clinics of North America 27, 269–281. 10.1016/s0889-857x(05)70201-5.

49. Gurvits, G.E., and Tan, A. (2013). Burning mouth syndrome. World journal of gastroenterology 19, 665–672. 10.3748/wjg.v19.i5.665.

50. Samaranayake, L.P., Lamb, A.B., Lamey, P.J., and MacFarlane, T.W. (1989). Oral carriage of Candida species and coliforms in patients with burning mouth syndrome. Journal of oral pathology & medicine : official publication of the International Association of Oral Pathologists and the American Academy of Oral Pathology 18, 233–235. 10.1111/j.1600-0714.1989.tb00769.x.

51. Lövgren, A., Häggman-Henrikson, B., Visscher, C.M., Lobbezoo, F., Marklund, S., and Wänman, A. (2016). Temporomandibular pain and jaw dysfunction at different ages covering the lifespan--A population based study. European journal of pain (London, England) 20, 532–540. 10.1002/ejp.755.

52. Campello, C.P., Lima, E.L.S., Fernandes, R.S.M., Porto, M., and Muniz, M.T.C. (2022). TNF-α levels and presence of SNP-308G/A of TNF-α gene in temporomandibular disorder patients. Dental press journal of orthodontics 27, e2220159. 10.1590/2177-6709.27.1.e2220159.oar.

53. Sharma, S., Gupta, D.S., Pal, U.S., and Jurel, S.K. (2011). Etiological factors of temporomandibular joint disorders. National journal of maxillofacial surgery 2, 116–119. 10.4103/0975-5950.94463.

54. Scott, D.L., Wolfe, F., and Huizinga, T.W. (2010). Rheumatoid arthritis. Lancet (London, England) 376, 1094–1108. 10.1016/s0140-6736(10)60826-4.

55. Kroese, J.M., Volgenant, C.M.C., Crielaard, W., Loos, B., van Schaardenburg, D., Visscher, C.M., and Lobbezoo, F. (2021). Temporomandibular disorders in patients with early rheumatoid arthritis and at-risk individuals in the Dutch population: a cross-sectional study. RMD open 7. 10.1136/rmdopen-2020-001485.

56. Barut, K., Adrovic, A., Şahin, S., and Kasapçopur, Ö. (2017). Juvenile Idiopathic Arthritis. Balkan medical journal 34, 90–101. 10.4274/balkanmedj.2017.0111.

57. Larheim, T.A., and Haanaes, H.R. (1981). Micrognathia, temporomandibular joint changes and dental occlusion in juvenile rheumatoid arthritis of adolescents and adults. Scandinavian journal of dental research 89, 329–338. 10.1111/j.1600-0722.1981.tb01690.x.

58. Research, N.I.o.D.a.C. (2023). TMD (Temporomandibular Disorders). https://www.nidcr.nih.gov/health-info/tmd.

59. Bycroft, C., Freeman, C., Petkova, D., Band, G., Elliott, L.T., Sharp, K., Motyer, A., Vukcevic, D., Delaneau, O., O’Connell, J., et al. (2018). The UK Biobank resource with deep phenotyping and genomic data. Nature 562, 203–209. 10.1038/s41586-018-0579-z.

60. Chen, R., Davydov, E.V., Sirota, M., and Butte, A.J. (2010). Non-synonymous and synonymous coding SNPs show similar likelihood and effect size of human disease association. PloS one 5, e13574. 10.1371/journal.pone.0013574.

61. Sauna, Z.E., and Kimchi-Sarfaty, C. (2013). Synonymous mutations as a cause of human genetic disease. eLS. 10.1002/9780470015902.a0025173.

62. Hunt, R.C., Simhadri, V.L., Iandoli, M., Sauna, Z.E., and Kimchi-Sarfaty, C. (2014). Exposing synonymous mutations. Trends in Genetics 30, 308–321. 10.1016/j.tig.2014.04.006.

63. Spencer, P.S., Siller, E., Anderson, J.F., and Barral, J.M. (2012). Silent substitutions predictably alter translation elongation rates and protein folding efficiencies. Journal of molecular biology 422, 328–335. 10.1016/j.jmb.2012.06.010.

64. Schiffman, E., Ohrbach, R., Truelove, E., Look, J., Anderson, G., Goulet, J.P., List, T., Svensson, P., Gonzalez, Y., Lobbezoo, F., et al. (2014). Diagnostic Criteria for Temporomandibular Disorders (DC/TMD) for Clinical and Research Applications: recommendations of the International RDC/TMD Consortium Network* and Orofacial Pain Special Interest Group†. Journal of oral & facial pain and headache 28, 6–27. 10.11607/jop.1151.

65. Ferrillo, M., Giudice, A., Marotta, N., Fortunato, F., Di Venere, D., Ammendolia, A., Fiore, P., and de Sire, A. (2022). Pain Management and Rehabilitation for Central Sensitization in Temporomandibular Disorders: A Comprehensive Review. International journal of molecular sciences 23. 10.3390/ijms232012164.

66. Farré-Guasch, E., Aliberas, J.T., Spada, N.F., de Vries, R., Schulten, E., and Lobbezoo, F. (2023). The role of inflammatory markers in Temporomandibular Myalgia: A systematic review. The Japanese dental science review 59, 281–288. 10.1016/j.jdsr.2023.08.006.

67. Kim, S.-J., Park, Y.-H., Hong, S.-P., Cho, B.-O., Park, J.-W., and Kim, S.-G. (2003). The presence of bacteria in the synovial fluid of the temporomandibular joint and clinical significance: preliminary study. Journal of Oral and Maxillofacial Surgery 61, 1156–1161. 10.1016/S0278-2391(03)00674-8.

68. Henry, C.H., Hughes, C.V., Gérard, H.C., Hudson, A.P., and Wolford, L.M. (2000). Reactive arthritis: preliminary microbiologic analysis of the human temporomandibular joint. Journal of oral and maxillofacial surgery : official journal of the American Association of Oral and Maxillofacial Surgeons 58, 1137-1142; discussion 1143-1134. 10.1053/joms.2000.9575.

69. Kim, Y., Gumpper, R.H., Liu, Y., Kocak, D.D., Xiong, Y., Cao, C., Deng, Z., Krumm, B.E., Jain, M.K., Zhang, S., et al. (2024). Bitter taste receptor activation by cholesterol and an intracellular tastant. Nature 628, 664–671. 10.1038/s41586-024-07253-y.

70. Medapati, M.R., Singh, N., Bhagirath, A.Y., Duan, K., Triggs-Raine, B., Batista, E.L., Jr., and Chelikani, P. (2021). Bitter taste receptor T2R14 detects quorum sensing molecules from cariogenic Streptococcus mutans and mediates innate immune responses in gingival epithelial cells. FASEB journal : official publication of the Federation of American Societies for Experimental Biology 35, e21375. 10.1096/fj.202000208R.

71. Medapati, M.R., Bhagirath, A.Y., Singh, N., Schroth, R.J., Bhullar, R.P., Duan, K., and Chelikani, P. (2021). Bitter Taste Receptor T2R14 Modulates Gram-Positive Bacterial Internalization and Survival in Gingival Epithelial Cells. International journal of molecular sciences 22. 10.3390/ijms22189920.

72. Kaur, K., Turner, A., Jones, P., Sculley, D., Veysey, M., Lucock, M., Wallace, J., and Beckett, E.L. (2021). A Cross-Sectional Study of Bitter-Taste Receptor Genotypes, Oral Health, and Markers of Oral Inflammation. Oral 1, 122–138.

73. Locker, D., and Miller, Y. (1994). Evaluation of subjective oral health status indicators. J Public Health Dent 54, 167–176. 10.1111/j.1752-7325.1994.tb01209.x.

74. Suresh, K., and Chandrashekara, S. (2012). Sample size estimation and power analysis for clinical research studies. Journal of human reproductive sciences 5, 7–13. 10.4103/0974-1208.97779.

75. Purcell, S., Neale, B., Todd-Brown, K., Thomas, L., Ferreira, M.A., Bender, D., Maller, J., Sklar, P., de Bakker, P.I., Daly, M.J., and Sham, P.C. (2007). PLINK: a tool set for whole-genome association and population-based linkage analyses. American journal of human genetics 81, 559–575. 10.1086/519795.

76. Team, R.C. (2021). R: A language and environment for statistical computing. R Foundation for Statistical Computing, Vienna, Austria. https://www.R-project.org/.

77. Sayers, E.W., Bolton, E.E., Brister, J.R., Canese, K., Chan, J., Comeau, D.C., Connor, R., Funk, K., Kelly, C., Kim, S., et al. (2022). Database resources of the national center for biotechnology information. Nucleic Acids Res 50, D20–d26. 10.1093/nar/gkab1112.

78. McLaren, W., Gil, L., Hunt, S.E., Riat, H.S., Ritchie, G.R.S., Thormann, A., Flicek, P., and Cunningham, F. (2016). The Ensembl Variant Effect Predictor. Genome Biology 17, 122. 10.1186/s13059-016-0974-4.

79. Yang, J., and Zhang, Y. (2015). I-TASSER server: new development for protein structure and function predictions. Nucleic Acids Res 43, W174–181. 10.1093/nar/gkv342.

80. Han, X., Steven, K., Qassim, A., Marshall, H.N., Bean, C., Tremeer, M., An, J., Siggs, O.M., Gharahkhani, P., Craig, J.E., et al. (2021). Automated AI labeling of optic nerve head enables insights into cross-ancestry glaucoma risk and genetic discovery in >280,000 images from UKB and CLSA. The American Journal of Human Genetics 108, 1204–1216. 10.1016/j.ajhg.2021.05.005.

81. Bassim, C.W., MacEntee, M.I., Nazmul, S., Bedard, C., Liu, S., Ma, J., Griffith, L.E., and Raina, P. (2020). Self-reported oral health at baseline of the Canadian Longitudinal Study on Aging. Community dentistry and oral epidemiology 48, 72–80. 10.1111/cdoe.12506.

82. Auton, A., Abecasis, G.R., Altshuler, D.M., Durbin, R.M., Abecasis, G.R., Bentley, D.R., Chakravarti, A., Clark, A.G., Donnelly, P., Eichler, E.E., et al. (2015). A global reference for human genetic variation. Nature 526, 68–74. 10.1038/nature15393.

83. Wickham, H. (2016). ggplot2: Elegant Graphics for Data Analysis. Springer-Verlag New York. https://ggplot2.tidyverse.org.

84. Akoglu, H. (2018). User’s guide to correlation coefficients. Turkish journal of emergency medicine 18, 91–93. 10.1016/j.tjem.2018.08.001.

85. (RTH), C.f.N.-c.R.i.T.a.H. RNAsnp: Predicting RNA secondary structure changes upon SNPs. https://rth.dk/resources/rnasnp/.

