## Supplementary material for "Bitter taste genetics and oral health in Canadian Longitudinal Study on Aging": Document S1

**Table S1. Allele distributions of *TAS2R* SNPs in the CLSA and 1KGP, Related to Figure 1.** Data are presented as minor allele frequencies (MAF) in CLSA European ethnicity and corresponding allele frequencies (AF) in 1KGP European population.

| rs ID | SNP[a] | MA[b] | Gene | Consequence | Amino Acid Change | CLSA MAF | Allele Count[c] | 1KGP AF | Adjusted p-value[d] |
|---|---|---|---|---|---|---|---|---|---|
| rs41466 | chr5:9627670:C:T | C | *TAS2R1* | 3 prime UTR | - | 0.45 | 45198 | 0.44 | 10.814 |
| rs41467 | chr5:9627977:A:C | A | *TAS2R1* | 3 prime UTR | - | 0.45 | 45862 | 0.44 | 20.777 |
| rs41468 | chr5:9627996:G:A | G | *TAS2R1* | 3 prime UTR | - | 0.46 | 45202 | 0.44 | 10.239 |
| rs2234235 | chr5:9629183:A:G | G | *TAS2R1* | Synonymous | Leu 284 | 0.03 | 45942 | 0.05 | **0.021** |
| rs2234233 | chr5:9629417:G:A | A | *TAS2R1* | Missense | Arg 206 Trp | 0.17 | 45934 | 0.15 | 0.919 |
| rs138940415 | chr5:9630121:G:A | A | *TAS2R1* | 5 prime UTR | - | 0.02 | 45600 | 0.02 | 27.966 |
| rs847923 | chr7:12491252:G:C | C | *TAS2R2P* | NA | - | 0.31 | 45764 | 0.30 | 20.456 |
| rs35620026 | chr7:12491542:C:T | T | *TAS2R2P* | NA | - | 0.12 | 45798 | 0.13 | 24.972 |
| rs6963925 | chr7:12491554:C:T | T | *TAS2R2P* | NA | - | 0.26 | 45734 | 0.26 | 34.886 |
| rs12535592 | chr7:12491615:A:G | G | *TAS2R2P* | NA | - | 0.25 | 45814 | 0.25 | 34.600 |
| rs73058722 | chr7:12491991:A:G | G | *TAS2R2P* | NA | - | 0.25 | 45820 | 0.25 | 35.000 |
| rs765007 | chr7:141764114:T:C | C | *TAS2R3* | 5 prime UTR | - | 0.50 | 45848 | 0.47 | 0.277 |
| rs2270009 | chr7:141764965:C:T | T | *TAS2R3* | Synonymous | Gly 269 | 0.49 | 45864 | 0.46 | 0.349 |
| rs33920115 | chr7:141777387:G:A | A | *TAS2R4* | 5 prime UTR | - | 0.50 | 45912 | 0.47 | 0.300 |
| rs2233998 | chr7:141778508:T:C | C | *TAS2R4* | Missense | Phe 7 Ser | 0.50 | 45944 | 0.47 | 0.277 |
| rs2234001 | chr7:141778774:G:C | G | *TAS2R4* | Missense | Leu 96 Val | 0.50 | 45910 | 0.53 | 0.472 |
| rs2234002 | chr7:141779000:G:A | G | *TAS2R4* | Missense | Asn 171 Ser | 0.50 | 45944 | 0.53 | 0.449 |
| rs57641758 | chr7:141779930:A:G | A | *TAS2R4* | 3 prime UTR | - | 0.50 | 45894 | 0.53 | 0.427 |
| rs139145080 | chr7:141779944:A:G | G | *TAS2R4* | 3 prime UTR | - | 0.02 | 45912 | 0.02 | 9.225 |
| rs150967230 | chr7:141780098:C:T | T | *TAS2R4* | 3 prime UTR | - | 0.01 | 45794 | 0.02 | 0.184 |
| rs35127278 | chr7:141780293:G:T | T | *TAS2R4* | 3 prime UTR | - | 0.50 | 45908 | 0.47 | 0.307 |
| rs2214838 | chr7:141780413:A:G | G | *TAS2R4* | 3 prime UTR | - | 0.50 | 45906 | 0.47 | 0.307 |
| rs2190243 | chr7:141780676:C:G | C | *TAS2R4* | 3 prime UTR | - | 0.50 | 45898 | 0.53 | 0.483 |
| rs2190244 | chr7:141780772:A:G | G | *TAS2R4* | 3 prime UTR | - | 0.50 | 45908 | 0.47 | 0.307 |
| rs17464086 | chr7:141781136:G:A | G | *TAS2R4* | 3 prime UTR | - | 0.50 | 45880 | 0.53 | 0.508 |
| rs2234009 | chr7:141790231:C:T | T | *TAS2R5* | 5 prime UTR | - | 0.03 | 45948 | 0.04 | 0.058 |
| rs2234010 | chr7:141790232:G:A | A | *TAS2R5* | 5 prime UTR | - | 0.03 | 45948 | 0.02 | 0.854 |
| rs2234012 | chr7:141790307:A:G | G | *TAS2R5* | 5 prime UTR | - | 0.50 | 45878 | 0.47 | 0.406 |
| rs2227264 | chr7:141790438:G:T | T | *TAS2R5* | Missense | Ser 26 Ile | 0.50 | 45868 | 0.47 | 0.332 |
| rs117638090 | chr7:141787930:G:A | A | *TAS2R6P* | Upstream gene | - | 0.02 | 45912 | 0.02 | 9.060 |
| rs1859645 | chr7:141788088:A:G | G | *TAS2R6P* | Upstream gene | - | 0.50 | 45882 | 0.47 | 0.417 |
| rs11761380 | chr7:141788474:A:C | C | *TAS2R6P* | Upstream gene | - | 0.50 | 45882 | 0.47 | 0.417 |
| rs619381 | chr12:10801659:C:T | T | *TAS2R7* | Missense | Met 304 Ile | 0.12 | 45924 | 0.09 | <0.0001 |
| rs139604652 | chr12:10801984:A:T | T | *TAS2R7* | Missense | Val 196 Glu | 0.01 | 45606 | 0.01 | 33.554 |
| rs143804727 | chr12:10806297:T:G | G | *TAS2R8* | Missense | Glu 228 Asp | 0.02 | 45936 | 0.01 | 0.319 |
| rs1548803 | chr12:10806432:C:T | C | *TAS2R8* | Synonymous | Leu 183 | 0.41 | 45766 | 0.32 | <0.0001 |
| rs3741845 | chr12:10809516:A:G | A | *TAS2R9* | Missense | Ala 187 Val | 0.40 | 45938 | 0.32 | <0.0001 |
| rs142359564 | chr12:10895096:C:T | T | *TAS2R12P* | NA | - | 0.01 | 45908 | 0.01 | 12.386 |
| rs16925338 | chr12:10895290:T:A | A | *TAS2R12P* | NA | - | 0.02 | 45640 | 0.01 | 10.713 |
| rs11054065 | chr12:10895780:G:A | A | *TAS2R12P* | NA | - | 0.02 | 45946 | 0.01 | **0.023** |
| rs1015442 | chr12:10908096:T:C | T | *TAS2R13* | 3 prime UTR | - | 0.43 | 45930 | 0.38 | **0.000** |
| rs1015443 | chr12:10908523:T:C | T | *TAS2R13* | Missense | Ser 259 Asn | 0.43 | 45938 | 0.38 | **0.000** |
| rs3851584 | chr12:10937478:G:T | G | *TAS2R14* | 3 prime UTR | - | 0.43 | 45904 | 0.38 | **0.000** |

| rs ID | SNP[a] | MA[b] | Gene | Consequence | Amino Acid Change | CLSA MAF | Allele Count[c] | 1KGP AF | Adjusted p-value[d] |
|---|---|---|---|---|---|---|---|---|---|
| rs3851585 | chr12:10937794:C:G | G | *TAS2R14* | 3 prime UTR | - | 0.02 | 45942 | 0.03 | **0.010** |
| rs3936285 | chr12:10937839:T:A | A | *TAS2R14* | 3 prime UTR | - | 0.02 | 45942 | 0.03 | **0.010** |
| rs35804287 | chr12:10938607:G:A | A | *TAS2R14* | Missense | Leu 201 Phe | 0.02 | 45942 | 0.03 | 0.468 |
| rs3741843 | chr12:10938833:C:T | C | *TAS2R14* | Synonymous | Arg 125 | 0.16 | 45912 | 0.14 | 0.074 |
| rs16925868 | chr12:10938952:T:C | C | *TAS2R14* | Missense | Thr 86 Ala | 0.02 | 45946 | 0.03 | **0.012** |
| rs7138535 | chr12:10939094:T:A | A | *TAS2R14* | Synonymous | Gly 38 | 0.25 | 45916 | 0.22 | **0.006** |
| rs11054092 | chr12:10964560:T:C | C | *TAS2R15P* | NA | - | 0.25 | 45838 | 0.22 | **0.007** |
| rs11054093 | chr12:10964615:G:C | C | *TAS2R15P* | NA | - | 0.25 | 45838 | 0.22 | **0.007** |
| rs11054094 | chr12:10964693:G:A | A | *TAS2R15P* | NA | - | 0.25 | 45838 | 0.22 | **0.007** |
| rs4763599 | chr12:10964722:A:G | G | *TAS2R15P* | NA | - | 0.25 | 45838 | 0.22 | **0.007** |
| rs17810798 | chr12:10964833:G:C | C | *TAS2R15P* | NA | - | 0.25 | 45838 | 0.22 | **0.007** |
| rs11054095 | chr12:10965032:G:C | C | *TAS2R15P* | NA | - | 0.25 | 45830 | 0.22 | **0.008** |
| rs11054096 | chr12:10965197:T:C | C | *TAS2R15P* | NA | - | 0.25 | 45798 | 0.22 | **0.007** |
| rs1204014 | chr7:122994789:C:T | T | *TAS2R16* | Synonymous | Thr 282 | 0.04 | 45754 | 0.04 | 8.615 |
| rs860170 | chr7:122994970:C:T | C | *TAS2R16* | Missense | His 222 Arg | 0.33 | 45946 | 0.31 | 11.462 |
| rs7310047 | chr12:11158921:G:A | G | *TAS2R18P* | NA | - | 0.48 | 45914 | 0.49 | 11.925 |
| rs7296270 | chr12:11158991:A:T | A | *TAS2R18P* | NA | - | 0.16 | 45894 | 0.13 | **0.011** |
| rs61928603 | chr12:11159188:C:T | C | *TAS2R18P* | NA | - | 0.18 | 45916 | 0.16 | 1.273 |
| rs61928604 | chr12:11159348:C:T | C | *TAS2R18P* | NA | - | 0.16 | 45926 | 0.13 | **0.018** |
| rs2290318 | chr12:11159359:G:C | C | *TAS2R18P* | NA | - | 0.32 | 45918 | 0.36 | **0.005** |
| rs2290319 | chr12:11159427:C:A | A | *TAS2R18P* | NA | - | 0.32 | 45916 | 0.36 | **0.005** |
| rs77837442 | chr12:11021687:C:T | T | *TAS2R19* | Stop-gained | Trp 295 * | 0.01 | 45918 | 0.01 | 15.228 |
| rs1868769 | chr12:11022154:G:A | G | *TAS2R19* | Synonymous | Leu 140 | 0.20 | 45770 | 0.19 | 4.040 |
| rs115193179 | chr12:11022246:T:G | G | *TAS2R19* | Missense | Lys 109 Thr | 0.01 | 45918 | 0.01 | 15.370 |
| rs61912291 | chr12:10996791:T:G | G | *TAS2R20* | 3 prime UTR | - | 0.24 | 45812 | 0.21 | 0.100 |
| rs117458236 | chr12:10996860:C:T | T | *TAS2R20* | 3 prime UTR | - | 0.02 | 45930 | 0.02 | 35.000 |
| rs1450839 | chr12:10996933:A:G | G | *TAS2R20* | 3 prime UTR | - | 0.33 | 45948 | 0.38 | **0.000** |
| rs10845279 | chr12:10997112:C:A | A | *TAS2R20* | Missense | Arg 255 Leu | 0.33 | 45948 | 0.38 | **0.000** |
| rs10845280 | chr12:10997121:A:G | G | *TAS2R20* | Missense | Phe 252 Ser | 0.33 | 45946 | 0.38 | **0.000** |
| rs10845281 | chr12:10997170:T:C | C | *TAS2R20* | Missense | Ile 236 Val | 0.33 | 45948 | 0.38 | **0.000** |
| rs12226919 | chr12:10997434:G:T | T | *TAS2R20* | Missense | His 148 Asn | 0.33 | 45948 | 0.38 | **0.000** |
| rs12226920 | chr12:10997447:G:T | T | *TAS2R20* | Missense | His 143 Gln | 0.33 | 45946 | 0.38 | **0.000** |
| rs79420812 | chr12:10997455:C:T | T | *TAS2R20* | Missense | Val 141 Ile | 0.19 | 45286 | 0.22 | **0.001** |
| rs11054142 | chr12:10997615:G:A | A | *TAS2R20* | Synonymous | Ala 87 | 0.33 | 45942 | 0.38 | **0.000** |
| rs7135018 | chr12:10997641:T:C | C | *TAS2R20* | Missense | Lys 79 Glu | 0.24 | 45904 | 0.21 | 0.098 |
| rs11054143 | chr12:10997720:T:C | C | *TAS2R20* | Synonymous | Ala 52 | 0.33 | 45942 | 0.38 | **0.000** |
| rs4388985 | chr12:10997952:G:A | G | *TAS2R20* | 5 prime UTR | - | 0.16 | 45900 | 0.14 | 0.090 |
| rs1463237 | chr12:10997980:C:T | C | *TAS2R20* | 5 prime UTR | - | 0.16 | 45884 | 0.14 | **0.049** |
| rs7301234 | chr12:10998285:G:A | A | *TAS2R20* | 5 prime UTR | - | 0.33 | 45938 | 0.38 | **0.000** |
| rs2600357 | chr12:11133155:G:A | G | *TAS2R30* | 3 prime UTR | - | 0.48 | 45902 | 0.49 | 12.082 |
| rs2600356 | chr12:11133203:G:T | G | *TAS2R30* | 3 prime UTR | - | 0.48 | 45902 | 0.49 | 12.082 |
| rs2599404 | chr12:11133489:A:C | A | *TAS2R30* | Missense | Leu 252 Phe | 0.48 | 45656 | 0.49 | 15.671 |
| rs112849631 | chr12:11133780:T:C | C | *TAS2R30* | Synonymous | Thr 155 | 0.02 | 45932 | 0.03 | **0.004** |
| rs116737741 | chr12:11030592:T:C | C | *TAS2R31* | Synonymous | Ser 248 | 0.15 | 45870 | 0.12 | **0.002** |
| rs10845294 | chr12:11030687:G:C | C | *TAS2R31* | Missense | Gln 217 Glu | 0.27 | 45796 | 0.27 | 22.105 |

| rs ID | SNP[a] | MA[b] | Gene | Consequence | Amino Acid Change | CLSA MAF | Allele Count[c] | 1KGP AF | Adjusted p-value[d] |
|---|---|---|---|---|---|---|---|---|---|
| rs10743938 | chr12:11030852:A:T | A | *TAS2R31* | Missense | Met 162 Leu | 0.20 | 45414 | 0.19 | 5.628 |
| rs10845295 | chr12:11031233:G:A | G | *TAS2R31* | Missense | Trp 35 Arg | 0.49 | 45826 | 0.52 | 0.615 |
| rs10246939 | chr7:141972804:T:C | C | *TAS2R38* | Missense | Ile 296 Val | 0.46 | 45922 | 0.46 | 26.426 |
| rs1726866 | chr7:141972905:G:A | G | *TAS2R38* | Missense | Val 262 Ala | 0.46 | 45916 | 0.46 | 26.896 |
| rs713598 | chr7:141973545:C:G | G | *TAS2R38* | Missense | Ala 49 Pro | 0.41 | 44464 | 0.42 | 10.330 |
| rs10260248 | chr7:143222638:C:A | A | *TAS2R40* | Missense | Ser 187 Tyr | 0.06 | 45946 | 0.06 | 29.364 |
| rs1404635 | chr7:143478061:G:A | A | *TAS2R41* | Synonymous | Thr 63 | 0.29 | 45896 | 0.25 | 0.096 |
| rs10278721 | chr7:143478252:C:T | T | *TAS2R41* | Missense | Pro 127 Leu | 0.29 | 45930 | 0.25 | 0.090 |
| rs76518630 | chr7:143478409:C:T | T | *TAS2R41* | Synonymous | Tyr 179 | 0.01 | 45746 | 0.02 | **0.035** |
| rs1650017 | chr12:11186007:G:C | C | *TAS2R42* | Missense | Pro 311 Ala | 0.16 | 45944 | 0.13 | **0.011** |
| rs1669411 | chr12:11186008:G:A | A | *TAS2R42* | Synonymous | Asn 310 | 0.16 | 45942 | 0.13 | **0.011** |
| rs1669412 | chr12:11186063:T:C | T | *TAS2R42* | Missense | Arg 292 Gln | 0.25 | 45948 | NA | NA |
| rs1451772 | chr12:11186144:C:T | C | *TAS2R42* | Missense | Tyr 265 Cys | 0.25 | 45942 | NA | NA |
| rs1669413 | chr12:11186175:A:C | C | *TAS2R42* | Missense | Trp 255 Gly | 0.16 | 45946 | 0.13 | **0.011** |
| rs5020531 | chr12:11186351:A:G | A | *TAS2R42* | Missense | Ser 196 Phe | 0.41 | 45946 | 0.35 | <0.0001 |
| rs1650019 | chr12:11186377:C:T | T | *TAS2R42* | Synonymous | Leu 187 | 0.16 | 45946 | 0.13 | **0.011** |
| rs35969491 | chr12:11186414:T:A | T | *TAS2R42* | Missense | Phe 175 Tyr | 0.41 | 45938 | 0.35 | <0.0001 |
| rs73260771 | chr12:11061761:C:T | T | *TAS2R46* | Synonymous | Thr 178 | 0.27 | 45920 | 0.28 | 20.458 |
| rs10772397 | chr12:10986084:C:T | C | *TAS2R50* | Synonymous | Pro 259 | 0.40 | 45946 | 0.34 | <0.0001 |
| rs1376251 | chr12:10986253:C:T | T | *TAS2R50* | Missense | Cys 203 Tyr | 0.32 | 45926 | 0.36 | **0.003** |
| rs66679979 | chr12:10986336:T:C | C | *TAS2R50* | Synonymous | Ser 175 | 0.15 | 45948 | 0.13 | **0.018** |
| rs61750008 | chr7:143443582:G:A | A | *TAS2R60* | Missense | Val 44 Met | 0.03 | 45938 | 0.04 | 0.254 |
| rs4595035 | chr7:143444382:T:C | T | *TAS2R60* | Synonymous | Arg 310 | 0.33 | 45936 | 0.34 | 11.534 |
| rs4726624 | chr7:143437150:G:A | A | *TAS2R62P* | NA | - | 0.13 | 45522 | 0.13 | 21.864 |
| rs34039200 | chr7:143437188:G:A | A | *TAS2R62P* | NA | - | 0.22 | 45936 | 0.23 | 4.662 |
| rs10239143 | chr7:143437699:C:G | G | *TAS2R62P* | NA | - | 0.13 | 45526 | 0.14 | 16.133 |
| rs6944279 | chr7:143437702:A:T | T | *TAS2R62P* | NA | - | 0.33 | 45330 | 0.34 | 10.871 |
| rs2708319 | chr12:11048442:A:G | A | *TAS2R63P* | NA | - | 0.49 | 45916 | 0.52 | 1.815 |
| rs35062230 | chr12:11048811:G:A | A | *TAS2R63P* | NA | - | 0.24 | 45944 | 0.21 | 0.102 |
| rs2597985 | chr12:11048888:T:C | T | *TAS2R63P* | NA | - | 0.49 | 45922 | 0.52 | 1.853 |
| rs2599394 | chr12:11049030:G:A | G | *TAS2R63P* | NA | - | 0.49 | 45712 | 0.52 | 1.327 |
| rs319268 | chr12:11179591:T:A | T | *TAS2R67P* | Downstream gene (SMIM10L1) | - | 0.25 | 45870 | NA | NA |
| rs73067307 | chr12:11179960:A:G | G | *TAS2R67P* | Downstream gene (SMIM10L1) | - | 0.03 | 45916 | 0.02 | 1.431 |
| rs319269 | chr12:11179986:A:C | C | *TAS2R67P* | Downstream gene (SMIM10L1) | - | 0.16 | 45910 | 0.13 | **0.011** |
| rs34648613 | chr12:11180118:T:A | T | *TAS2R67P* | Downstream gene (SMIM10L1) | - | 0.41 | 45864 | 0.35 | <0.0001 |
| rs184006834 | chr12:11180133:G:A | A | *TAS2R67P* | Downstream gene (SMIM10L1) | - | 0.04 | 45736 | 0.04 | 4.081 |

[a] Position in GRCh38 shown as chromosome:base pair position: reference allele: alternative allele
[b] Minor allele in CLSA
[c] Chi-squared test with Bonferroni correction. Significant values are bolded ($p < 0.05$).
[d] Non-missing allele count in the CLSA

**Table S2. Gene-wise count of variants with different allele distributions in CLSA compared to the 1KGP based on the adjusted chi-squared test results, Related to Figure 1.**

| *Gene* | Count of variants in CLSA | Count of variants with different allele distributions[a] | Proportion of variants with different allele distributions[b] |
|---|---|---|---|
| *TAS2R16* | 2 | 0 | 0.00 |
| *TAS2R19* | 3 | 0 | 0.00 |
| *TAS2R2P* | 5 | 0 | 0.00 |
| *TAS2R3* | 2 | 0 | 0.00 |
| *TAS2R38* | 3 | 0 | 0.00 |
| *TAS2R4* | 12 | 0 | 0.00 |
| *TAS2R40* | 1 | 0 | 0.00 |
| *TAS2R46* | 1 | 0 | 0.00 |
| *TAS2R5* | 4 | 0 | 0.00 |
| *TAS2R60* | 2 | 0 | 0.00 |
| *TAS2R62P* | 4 | 0 | 0.00 |
| *TAS2R63P* | 4 | 0 | 0.00 |
| *TAS2R6P* | 3 | 0 | 0.00 |
| *TAS2R7* | 2 | 0 | 0.00 |
| *TAS2R8* | 2 | 0 | 0.00 |
| *TAS2R9* | 1 | 0 | 0.00 |
| *TAS2R1* | 6 | 1 | 0.17 |
| *TAS2R67P* | 5 | 1 | 0.20 |
| *TAS2R30* | 4 | 1 | 0.25 |
| *TAS2R31* | 4 | 1 | 0.25 |
| *TAS2R12P* | 3 | 1 | 0.33 |
| *TAS2R41* | 3 | 1 | 0.33 |
| *TAS2R42* | 8 | 4 | 0.50 |
| *TAS2R50* | 3 | 2 | 0.67 |
| *TAS2R18P* | 6 | 4 | 0.67 |
| *TAS2R14* | 7 | 5 | 0.71 |
| *TAS2R20* | 15 | 11 | 0.73 |
| *TAS2R13* | 2 | 2 | 1.00 |
| *TAS2R15P* | 7 | 7 | 1.00 |

[a] The number of variants with significantly different allele distributions compared to the 1KGP.
[b] Count of variants with different allele distributions divided by the total count of variants in CLSA.

**Figure S1. Heatmaps showing the pairwise linkage disequilibrium (LD) between *TAS2R* variants, Related to Table 3.** A, B, and C represent the LD scores of variants on chromosomes 5, 7, and 12, respectively. The color intensity indicates the strength of LD, with $r^2$ values ranging from 0.2 to 1, while white cells represent $r^2 < 0.2$. Variants associated with sore jaw muscles are highlighted in red.

**A**

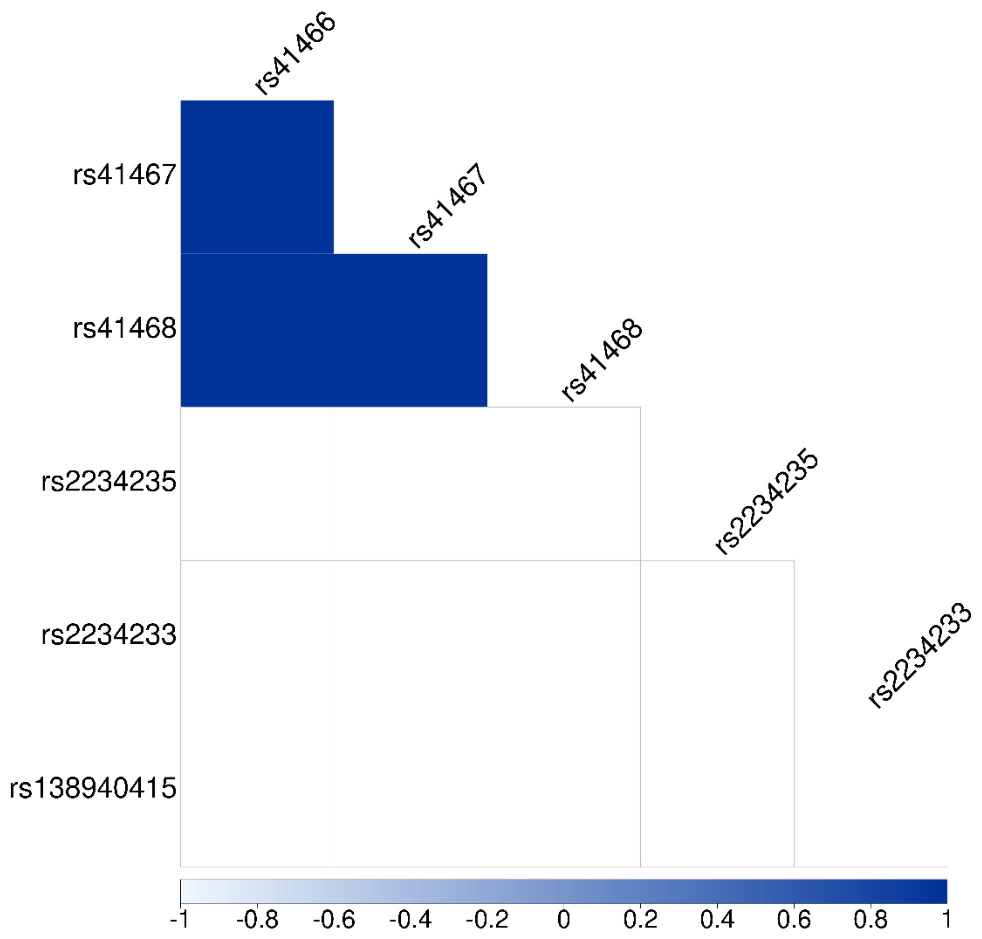

B

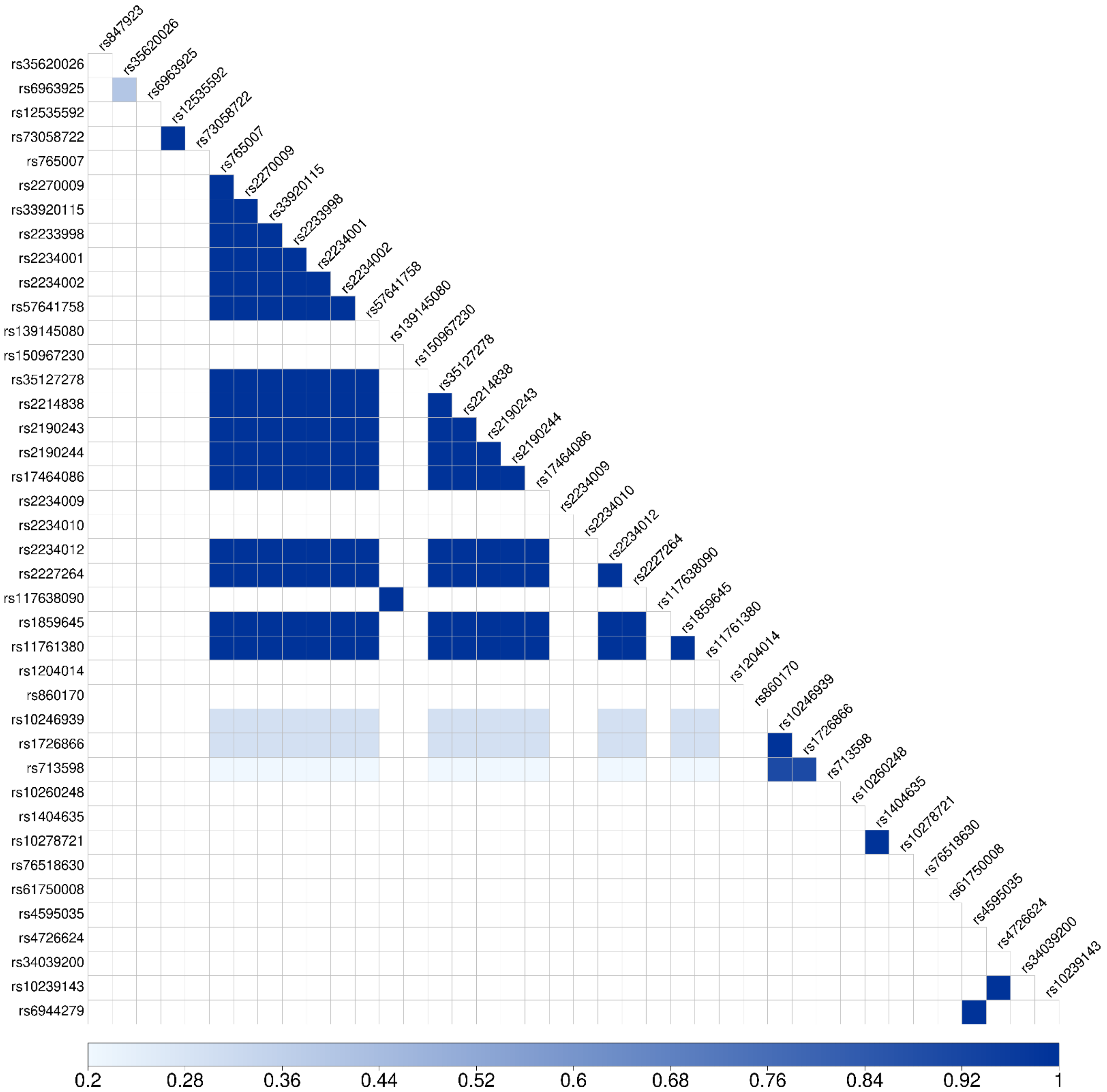
rs847923
rs35620026
rs6963925
rs12535592
rs73058722
rs765007
rs2270009
rs33920115
rs2233998
rs2234001
rs2234002
rs57641758
rs139145080
rs150967230
rs35127278
rs2214838
rs2190243
rs2190244
rs17464086
rs2234009
rs2234010
rs2234012
rs2227264
rs117638090
rs1859645
rs11761380
rs1204014
rs860170
rs10246939
rs1726866
rs713598
rs10260248
rs1404635
rs10278721
rs76518630
rs61750008
rs4595035
rs4726624
rs34039200
rs10239143
rs6944279
0.2
0.28
0.36
0.44
0.52
0.6
0.68
0.76
0.84
0.92
1

**C**

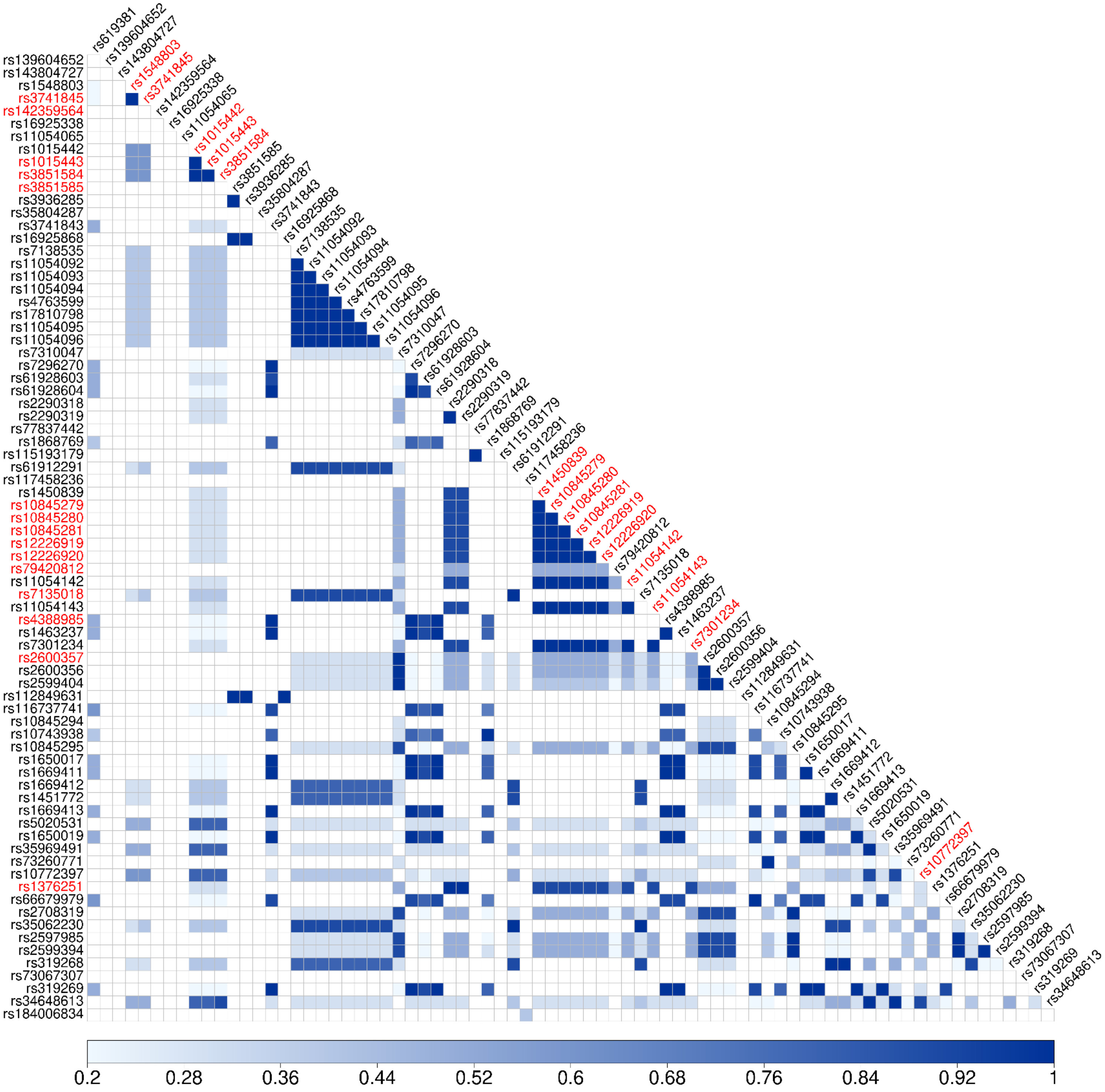

rs619381
rs139604652
rs143804727
rs1548803
rs3741845
rs142359564
rs16925338
rs11054065
rs1015442
rs1015443
rs3851584
rs3851585
rs3936285
rs35804287
rs3741843
rs16925868
rs7138535
rs11054092
rs11054093
rs11054094
rs4763599
rs17810798
rs11054095
rs11054096
rs7310047
rs7296270
rs61928603
rs61928604
rs2290318
rs2290319
rs77837442
rs1868769
rs115193179
rs61912291
rs117458236
rs1450839
rs10845279
rs10845280
rs10845281
rs12226919
rs12226920
rs79420812
rs11054142
rs7135018
rs11054143
rs4388985
rs1463237
rs7301234
rs2600357
rs2600356
rs2599404
rs112849631
rs116737741
rs10845294
rs10743938
rs10845295
rs1650017
rs1669411
rs1669412
rs1451772
rs1669413
rs5020531
rs1650019
rs35969491
rs73260771
rs10772397
rs1376251
rs66679979
rs2708319
rs35062230
rs2597985
rs2599394
rs319268
rs73067307
rs319269
rs34648613
rs184006834
0.2
0.28
0.36
0.44
0.52
0.6
0.68
0.76
0.84
0.92
1

**Figure S2. The optimal mRNA secondary structure and base pair probabilities predicted by RNAsnp, Related to Table 3.** (A-B) Seven variants that may impact mRNA secondary structure. (C-D) Eight variants that are not predicted to affect mRNA structure. The upper triangle of the matrices represents base pair probabilities for the wild-type local region, and the lower triangle represents those for the mutant sequence.

**A**

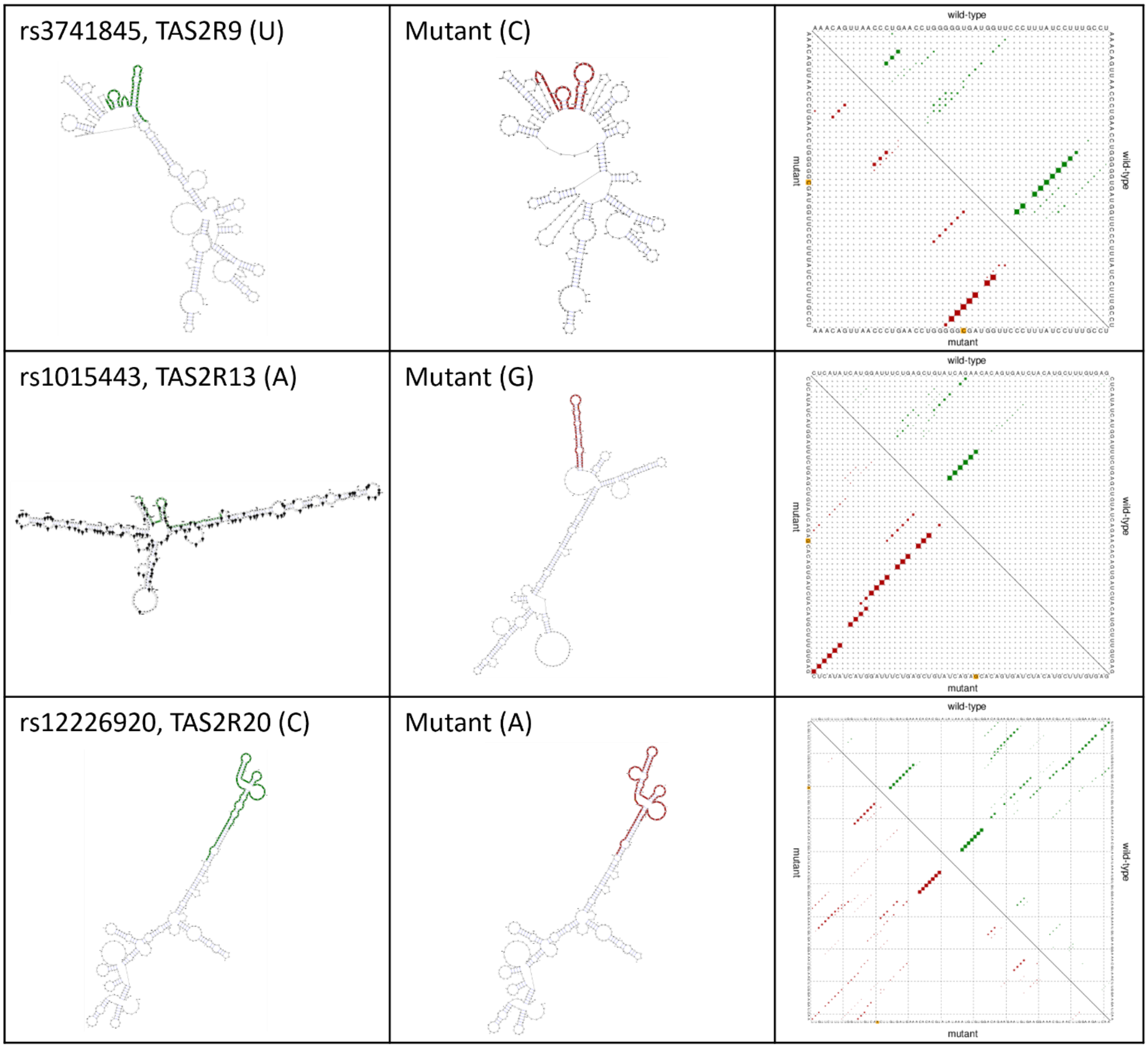

**B**

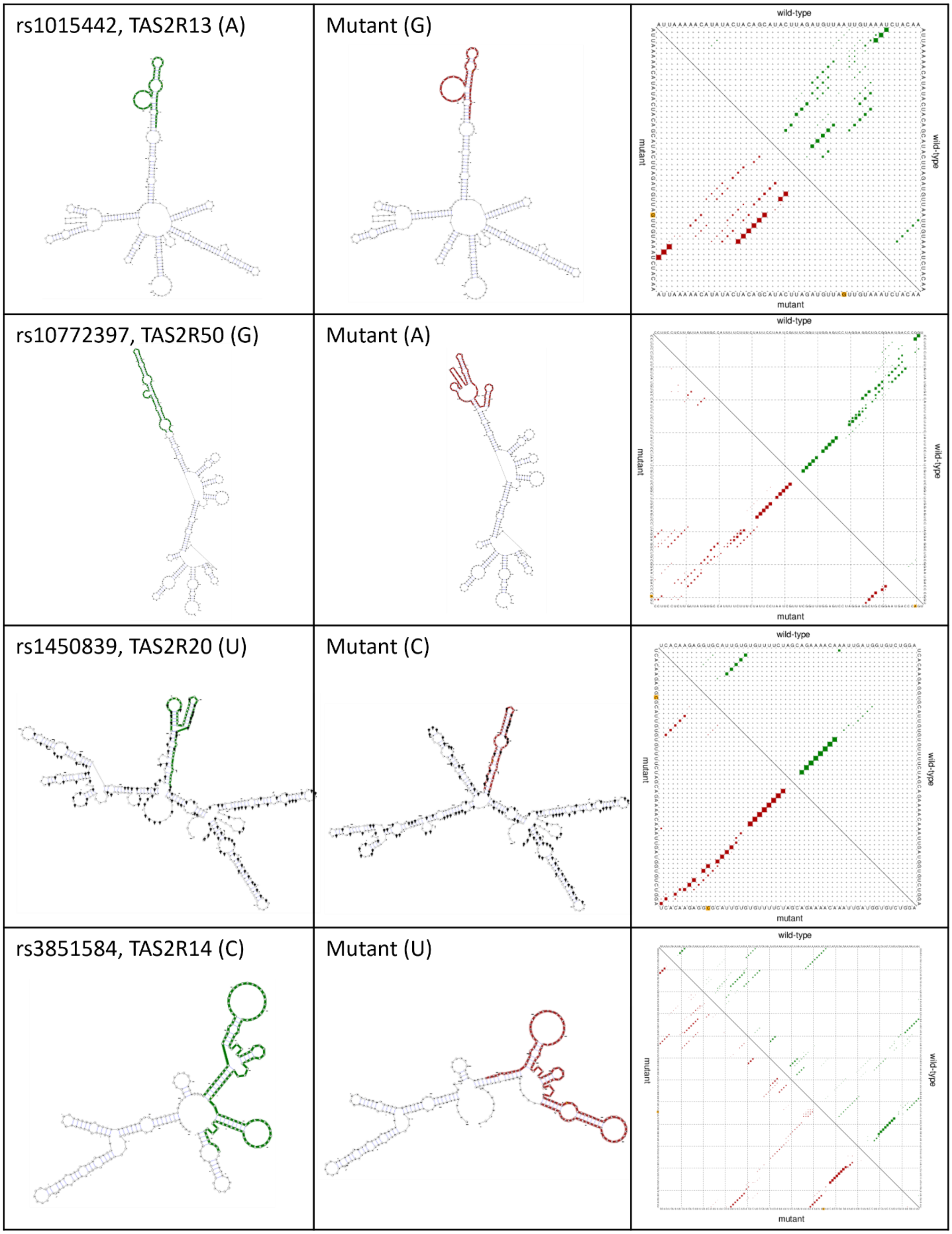
rs1015442, TAS2R13 (A)
Mutant (G)
wild-type
mutant
rs10772397, TAS2R50 (G)
Mutant (A)
wild-type
mutant
rs1450839, TAS2R20 (U)
Mutant (C)
wild-type
mutant
rs3851584, TAS2R14 (C)
Mutant (U)
wild-type
mutant

C

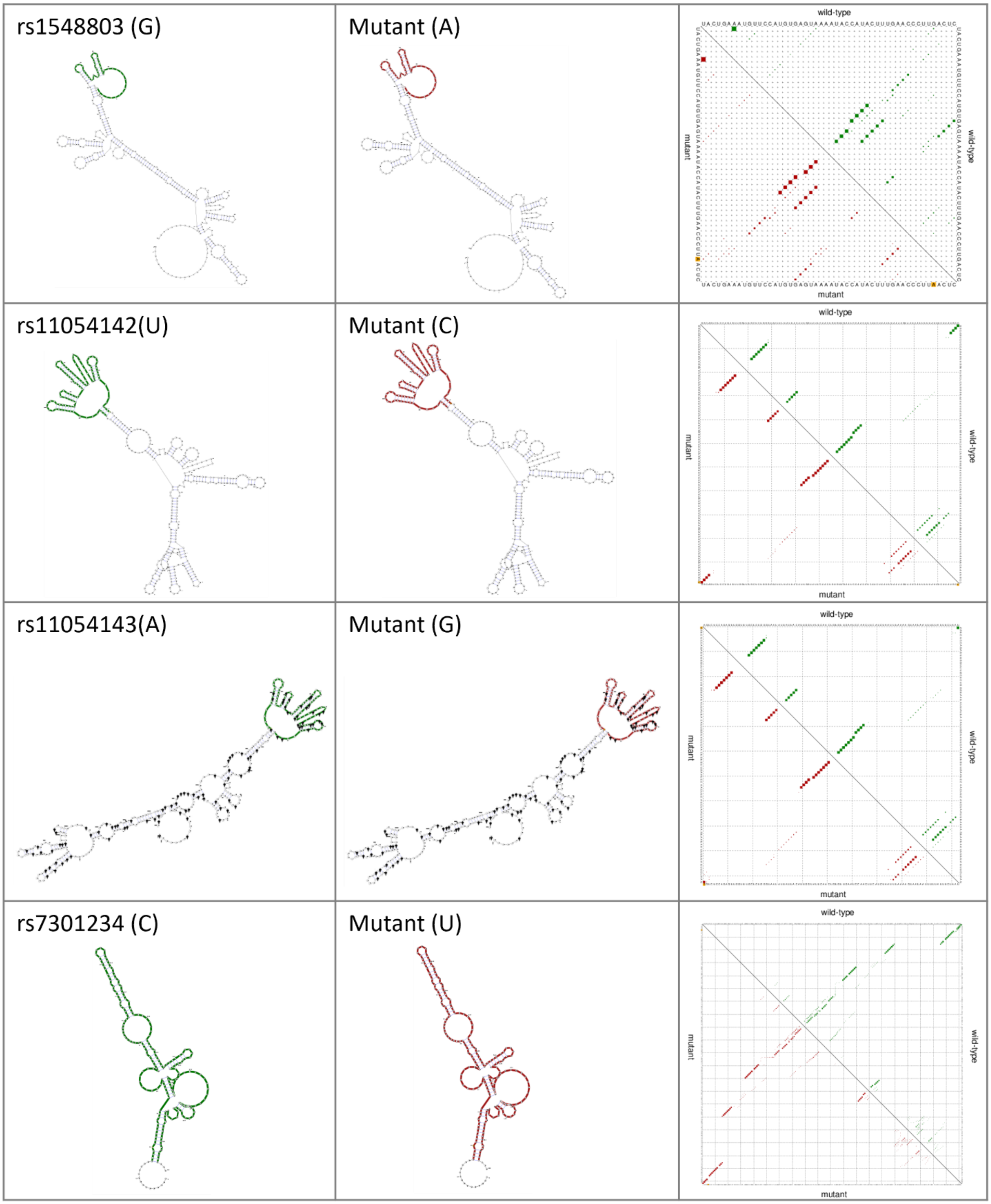
rs1548803 (G)
Mutant (A)
rs11054142(U)
Mutant (C)
rs11054143(A)
Mutant (G)
rs7301234 (C)
Mutant (U)
wild-type
mutant

**D**

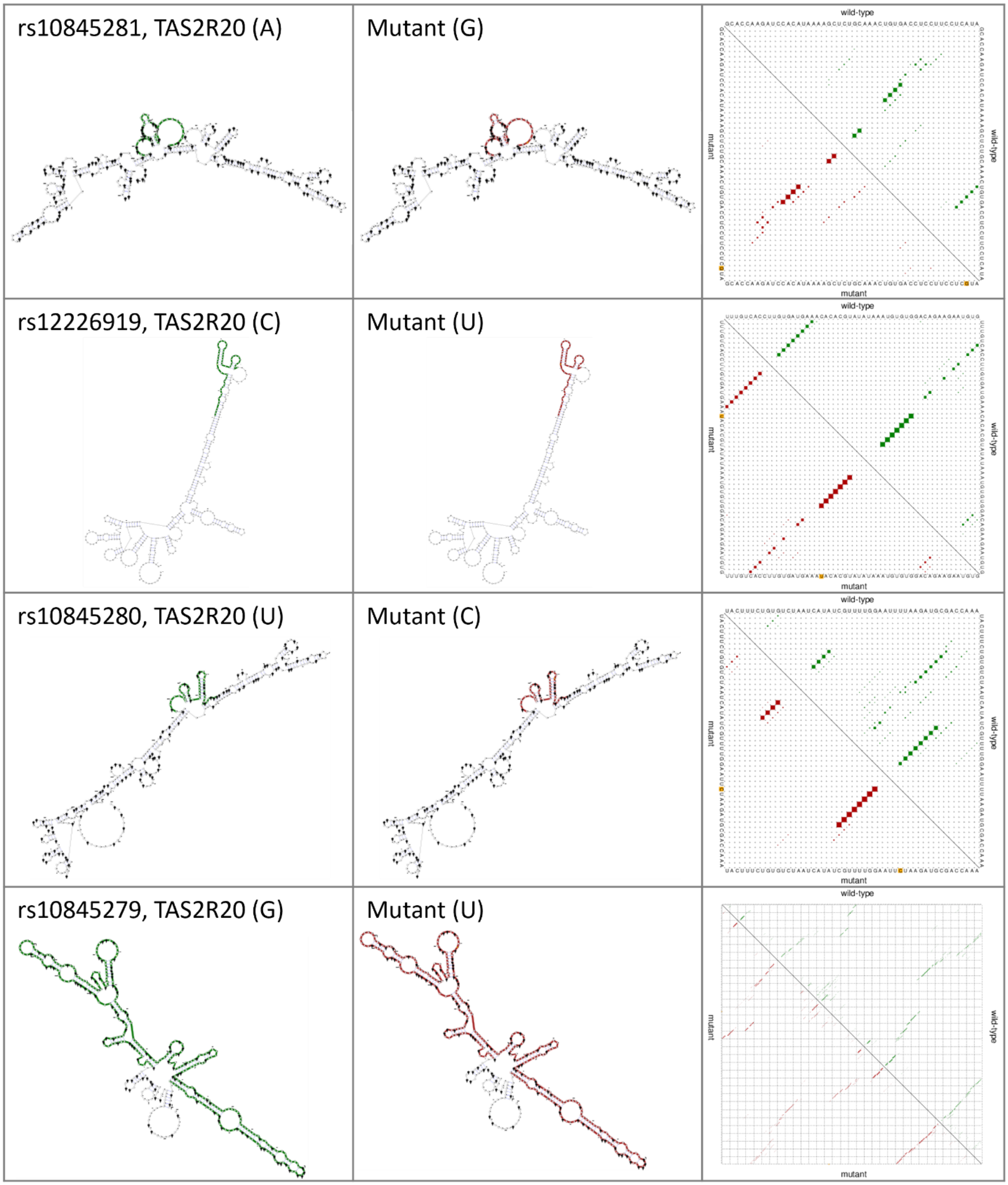
rs10845281, TAS2R20 (A)
Mutant (G)
rs12226919, TAS2R20 (C)
Mutant (U)
rs10845280, TAS2R20 (U)
Mutant (C)
rs10845279, TAS2R20 (G)
Mutant (U)
wild-type
mutant

**Figure S3. Manhattan plots displaying logistic regression results, Related to Figure 2.** The blue line denotes the significance threshold at (p = 0.05), while the red line marks the Bonferroni-corrected threshold.

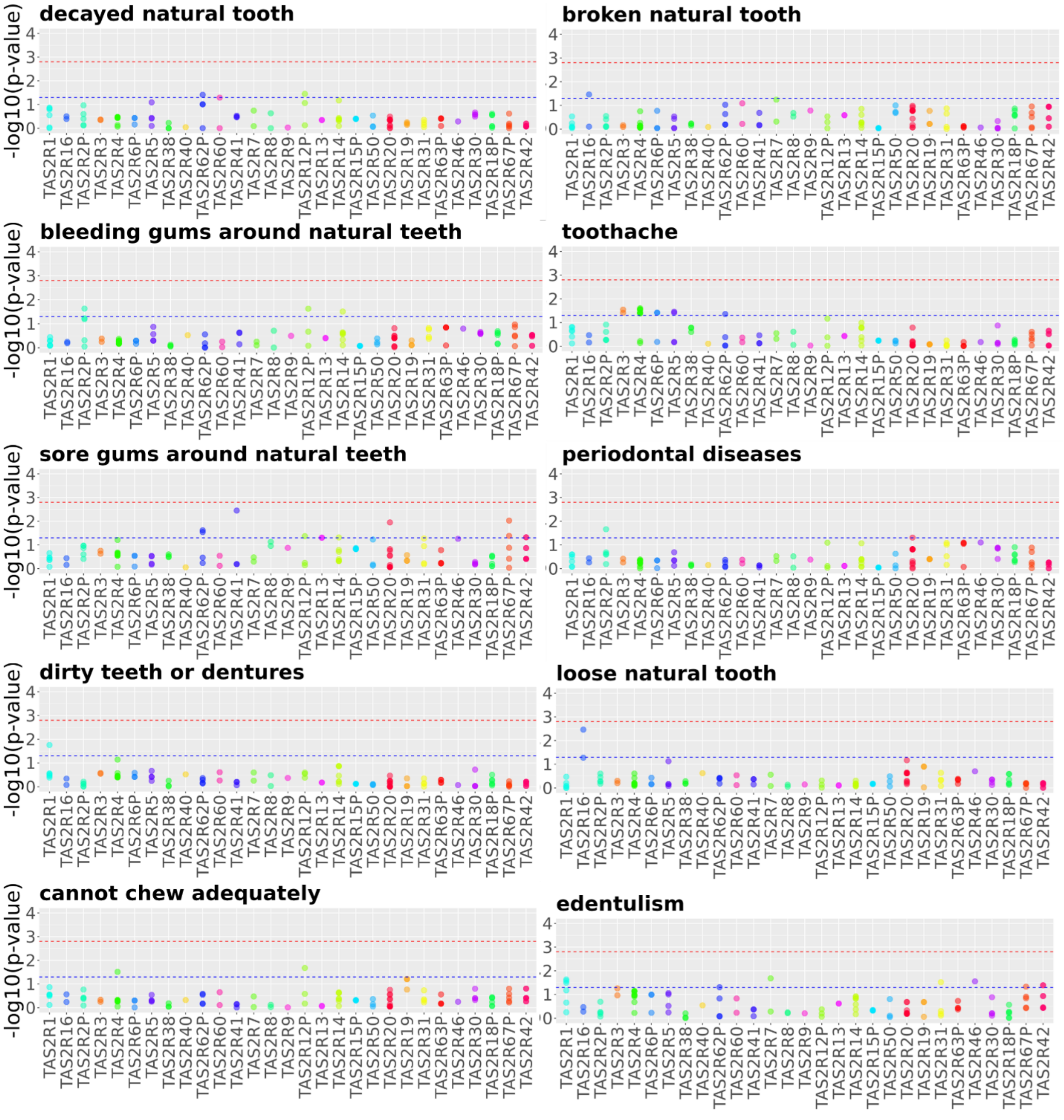

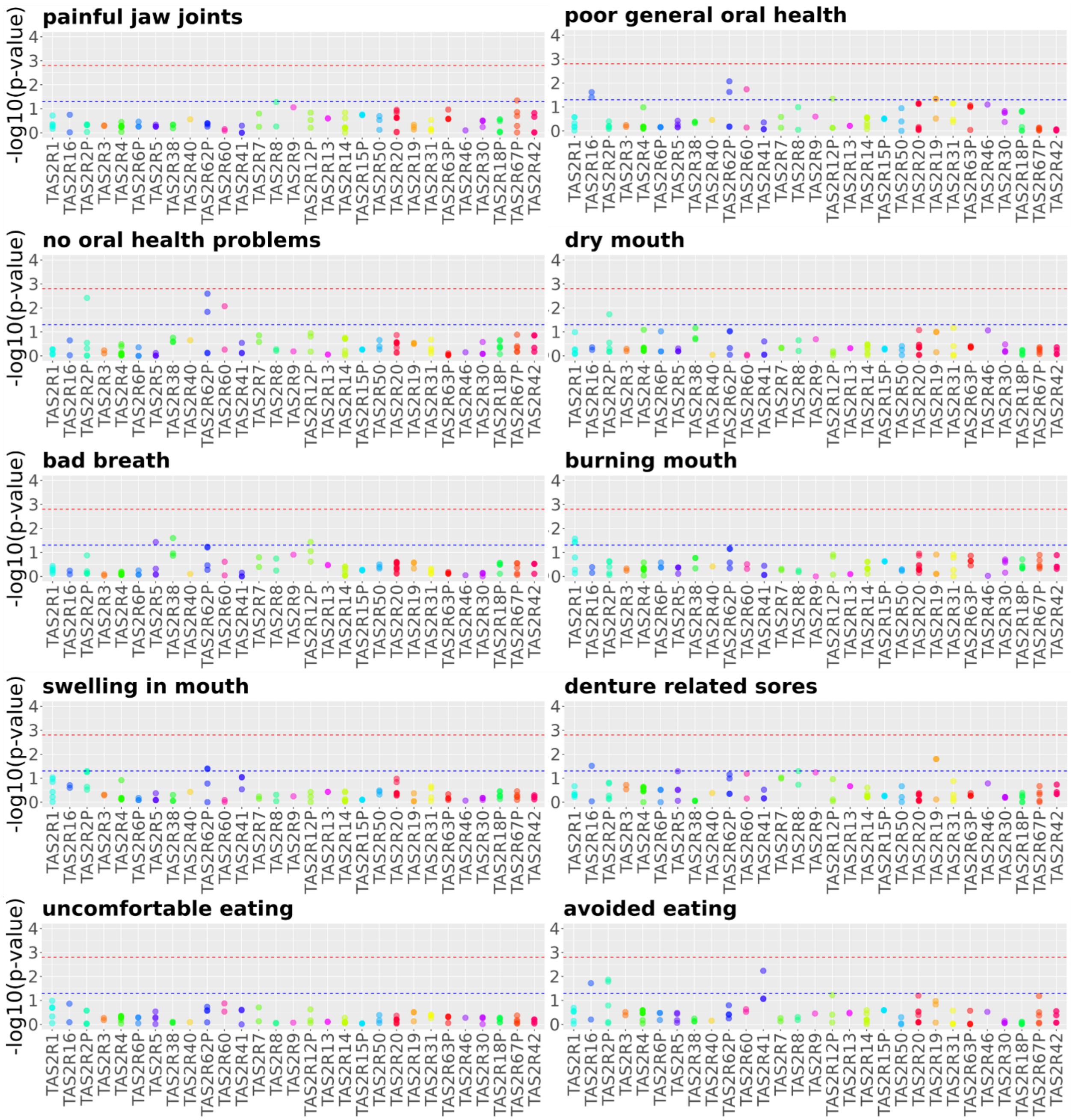
painful jaw joints
poor general oral health
no oral health problems
dry mouth
bad breath
burning mouth
swelling in mouth
denture related sores
uncomfortable eating
avoided eating
-log10(p-value)
0
1
2
3
4
TAS2R1 TAS2R16 TAS2R2P TAS2R3 TAS2R4 TAS2R6P TAS2R5 TAS2R38 TAS2R40 TAS2R62P TAS2R60 TAS2R41 TAS2R7 TAS2R8 TAS2R9 TAS2R12P TAS2R13 TAS2R14 TAS2R15P TAS2R50 TAS2R20 TAS2R19 TAS2R31 TAS2R63P TAS2R46 TAS2R30 TAS2R18P TAS2R67P TAS2R42

**Table S5. CLSA questionnaire for assessing oral health symptoms, Related to STAR Methods.**

| No. | Oral health symptoms | Answers | Question[a] |
|---|---|---|---|
| 1 | General oral health[b] | 1=excellent, 2=very good, 3=good, 4=fair, 5=poor | In general, would you say the health of your mouth is excellent, very good, good, fair, or poor? |
| 2 | Oral health problems - Toothache | 1=yes, 0=no | Have you experienced this in the past 12 months? |
| 3 | Oral health problems - Cannot chew adequately | 1=yes, 0=no | Have you experienced this in the past 12 months? |
| 4 | Oral health problems - Swelling in mouth | 1=yes, 0=no | Have you experienced this in the past 12 months? |
| 5 | Oral health problems - Dry mouth | 1=yes, 0=no | Have you experienced this in the past 12 months? |
| 6 | Oral health problems - Burning mouth | 1=yes, 0=no | Have you experienced this in the past 12 months? |
| 7 | Oral health problems - Sore jaw muscles | 1=yes, 0=no | Have you experienced this in the past 12 months? |
| 8 | Oral health problems - Painful jaw joints | 1=yes, 0=no | Have you experienced this in the past 12 months? |
| 9 | Oral health problems - Decayed natural tooth | 1=yes, 0=no | Have you experienced this in the past 12 months? |
| 10 | Oral health problems - Loose natural tooth | 1=yes, 0=no | Have you experienced this in the past 12 months? |
| 11 | Oral health problems - Broken natural tooth | 1=yes, 0=no | Have you experienced this in the past 12 months? |
| 12 | Oral health problems - Sore gums around natural teeth | 1=yes, 0=no | Have you experienced this in the past 12 months? |
| 13 | Oral health problems - Bleeding gums around natural teeth | 1=yes, 0=no | Have you experienced this in the past 12 months? |
| 14 | Oral health problems - Denture-related sores | 1=yes, 0=no | Have you experienced this in the past 12 months? |
| 15 | Oral health problems - Dirty teeth or dentures | 1=yes, 0=no | Have you experienced this in the past 12 months? |
| 16 | Oral health problems - Bad breath | 1=yes, 0=no | Have you experienced this in the past 12 months? |
| 17 | Oral health problems – No oral health problems | 1=yes, 0=no | Have you experienced this in the past 12 months? |
| 18 | Has one or more of the original teeth | 1=yes, 2=no | Do you have one or more of your original teeth? |
| 19 | Uncomfortable eating in last 12 months[b] | 1=often, 2=sometimes, 3= rarely, 4= never | In the past 12 months, how often have you found it uncomfortable to eat any food because of problems with your mouth? |
| 20 | Avoided eating in last 12 months[b] | 1=often, 2=sometimes, 3= rarely, 4= never | In the past 12 months, how often have you avoided eating particular foods because of problems with your mouth? |

[a] Questions asked from the participants for each of the variables.
[b] For statistical analyses, categorical variables with more than two values were dichotomized as follows: general oral health: good (excellent, very good, good) vs. poor (fair, poor); avoided eating and uncomfortable eating: yes (often, sometimes) vs. no (rarely, never).

**Figure S4. Correlation plot based on the Phi Coefficient (Mean Square Contingency Coefficient) for the binary phenotypes, Related to STAR Methods.**

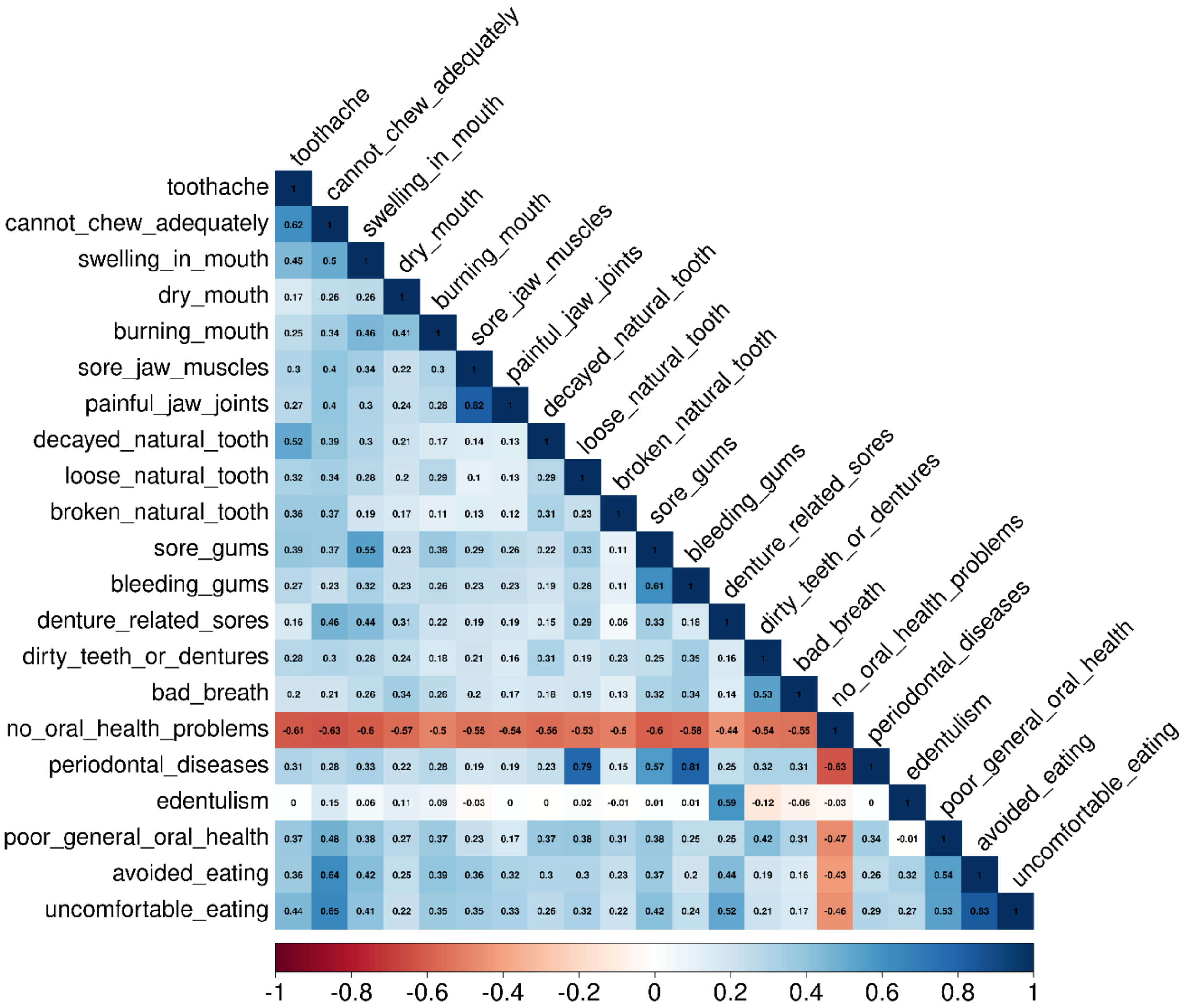

**Table S6. Association of variants linked to sore jaw muscles in the current study with rheumatoid arthritis, Related to STAR Methods.** The data is based on genome-wide association study (GWAS) results from the UK Biobank Imputed Version 3 dataset, generated by the Neale Lab.

| rs ID | Gene | MA | MAC[a] | Expected MAC[a] | Total allele count | beta | SE | P-value[b] |
|---|---|---|---|---|---|---|---|---|
| rs1548803 | *TAS2R8* | C | 4852.47 | 3313.85 | 424357 | 0.000774 | 0.000251 | **0.00202** |
| rs3741845 | *TAS2R9* | A | 4933 | 3238.03 | 431173 | 0.000806 | 0.000252 | **0.001361** |
| rs1015442 | *TAS2R13* | T | 4646.85 | 3502.98 | 407353 | 0.000665 | 0.000249 | **0.007495** |
| rs1015443 | *TAS2R13* | T | 4649 | 3503.51 | 407305 | 0.000679 | 0.000249 | **0.006297** |
| rs3851584 | *TAS2R14* | G | 4643.76 | 3507.59 | 406939 | 0.000674 | 0.000249 | **0.006744** |
| rs10772397 | *TAS2R50* | C | 4874 | 3283.05 | 427125 | 0.000717 | 0.000251 | **0.004253** |
| rs1450839 | *TAS2R20* | G | 2594 | 2556.37 | 229825 | 0.000232 | 0.000265 | 0.379467 |
| rs10845279 | *TAS2R20* | A | 2596 | 2557.2 | 229900 | 0.000241 | 0.000265 | 0.362629 |
| rs10845280 | *TAS2R20* | G | 2595 | 2557.28 | 229908 | 0.000233 | 0.000265 | 0.377791 |
| rs10845281 | *TAS2R20* | C | 2593 | 2556.32 | 229821 | 0.000226 | 0.000265 | 0.392459 |
| rs12226919 | *TAS2R20* | T | 2593 | 2554.69 | 229675 | 0.000237 | 0.000265 | 0.369795 |
| rs12226920 | *TAS2R20* | T | 2594 | 2556.36 | 229825 | 0.000233 | 0.000265 | 0.379405 |
| rs11054142 | *TAS2R20* | A | 2594 | 2556.36 | 229825 | 0.000233 | 0.000265 | 0.379405 |
| rs11054143 | *TAS2R20* | C | 2594 | 2556.42 | 229831 | 0.000232 | 0.000265 | 0.380261 |
| rs7301234 | *TAS2R20* | A | 2594 | 2556.42 | 229831 | 0.000232 | 0.000265 | 0.380261 |

MA: Minor allele
[a] Minor allele count in cases
[b] Statistically significant values are bolded.
